# Disentangled multimodal evolutionary representations for cross-virus predictive modeling of antigenic change

**DOI:** 10.64898/2026.02.12.705501

**Authors:** Zehong Zhang, Yuxi Lin, Siyun Zhong, Peng Zhou, Yixue Li, Fulong Yu

## Abstract

Antigenic evolution drives immune escape and complicates early assessment of emerging pathogens, especially when genomic surveillance is sparse and functional measurements lag behind real-world spread. Early prediction and generalization to unknown viruses require faithful and information-rich representations of viral evolution, a need that is not fully addressed by existing protein language model–based approaches. Here we introduce DERIVE, a flow-based generative framework that learns a disentangled latent representation of multimodal evolutionary history by integrating sequence homology with physicochemical and structural features. Using only pre-pandemic SARS-CoV-2 sequences, DERIVE prioritizes high-risk mutations, reconstructs strain-level evolutionary trajectories, forecasts immune escape and prevalence trends, and produces mutation-level effect maps. The learned representation transfers robustly across viral families, with strong concordance to functional data for influenza virus, HIV, rabies lyssavirus and chikungunya virus. Together, these results position DERIVE as a generalizable and interpretable framework for anticipating antigenic evolution and supporting early, actionable assessment of future “Disease X” threats.

## INTRODUCTION

Accurately anticipating how viruses evolve has become a central challenge for global health (*1, 2*). The rapid emergence of SARS-CoV-2 variants demonstrated how swiftly viral populations can shift, reshaping transmissibility and immune escape within weeks (*3-5*). During the earliest stages of an outbreak, when only sparse genome sequences and virtually no functional measurements are available, the ability to identify high-risk mutations or lineages and to forecast viral evolutionary trajectories before such variants rise in prevalence becomes critical for guiding vaccine updates, monoclonal antibody deployment and other public-health interventions.

Recent advances in representation learning suggest that such foresight may be attainable (*6-10*). Protein language models have shown that large-scale sequence corpora encode rich evolutionary constraints, and that latent representations learned from these data can recover functional structure even when measurements are sparse (*11-13*). However, antigenic evolution presents a qualitatively different challenge: natural viral sequences reflect only the aggregate fitness consequences of mutations rather than the specific biological pressures such as folding stability, receptor engagement, replication efficiency and immune escape that shape them (*14, 15*). These pressures operate along distinct and sometimes opposing axes of variation, meaning that their effects are fundamentally multidimensional. When these heterogeneous contributions are collapsed into a single fitness-proximal signal, their mechanistic meaning becomes obscured, limiting interpretability and restricting the ability to generalize predictive principles across antigens, lineages or viral families (*16, 17*).

Effective early forecasting therefore calls for representations that disentangle the constituent forces shaping viral evolution rather than merging them into a single axis. Such representations should allow phenotypes and other determinants to vary along distinct dimensions so that their contributions can be inferred without interference. Incorporating multimodal evolutionary histories, physicochemical descriptors and rapidly obtainable structural context further supports generalization across antigenically and structurally diverse viruses, an important consideration in early-stage Disease X scenarios (*18*).

Here we introduce DERIVE (DisEntanglement and Representation learning of antIgenic eVolutionary landscapEs), a unified flow-based framework that provides such a representation. DERIVE jointly models sequence homology, physicochemical signatures and structural history to learn a multimodal, disentangled latent space in which antigenicity is explicitly separated from other evolutionary pressures. Through this representation, DERIVE enables accurate mutation-level prediction, interpretable antigenic landscapes and evolutionary trajectories that generalize across viral families.

## RESULTS

### Overview of the DERIVE framework

DERIVE constructs a multimodal evolutionary history for each viral protein as the foundation for disentangled representation learning (Fig. 1A). We first assemble multiple sequence alignments (MSAs) from UniRef (*19*), MGnify (*20*) and BFD/Mgnfiy (*21, 22*) to capture sequence-level evolutionary information, given that MSAs provide the most broadly informative signal for modeling functional proteins (*23, 24*). A two-stage weighted sampling strategy is used to preserve both the depth and diversity of homologous sequences. Because protein–protein interactions, receptor engagement and immune recognition depend on physicochemical and structural properties that are not encoded in sequence alone, DERIVE augments the sequence history with complementary biophysical descriptors. These include curated AAindex clusters (*25*) emphasizing charge and hydrophobicity, structure-derived features capturing spatial neighborhood, disorder propensity (*26*), solvent exposure and coarse-grained force-field statistics (*27*), as well as an immunologically motivated indicator of N-linked glycosylation. Together, these components form a combined physicochemical– structural evolutionary history that extends the information available from MSAs and aligns with established principles in protein modeling and viral antigenicity.

**Fig. 1:**
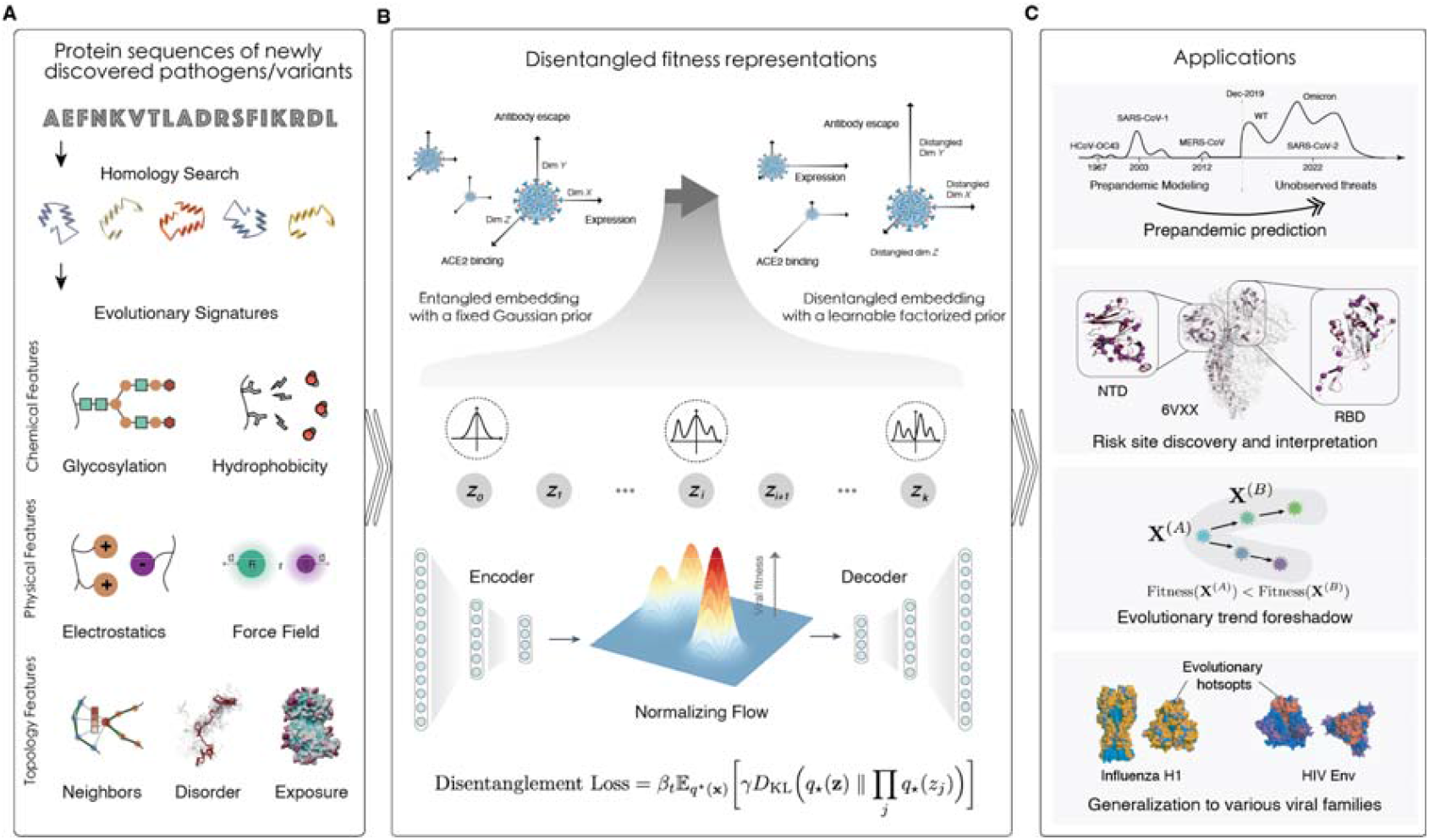
Overview of the DERIVE modeling and applications. (**A**) Construction of multimodal evolutionary histories. Each input protein sequence undergoes homology search against multiple sequence alignment (MSA) databases to obtain evolutionary relatives. These MSAs are augmented with amino-acid–level evolutionary signatures from three categories (**Methods**): chemical descriptors (N-linked glycosylation indicators and hydrophobicity indices), physical descriptors (electrostatics and coarse-grained force-field statistics) and structural–topological descriptors (residue-neighborhood graphs, disorder propensity and solvent exposure). (**B**) Disentanglement of latent evolutionary factors for multimodal evolutionary histories. DERIVE employs an encoder–decoder variational autoencoder equipped with a learnable normalizing-flow prior and a total-correlation regularization term (**Methods**). The flow prior reshapes the latent distribution away from an isotropic and coupled form toward a multimodal structure that separates distinct biological axes of variation. The resulting disentangled latent space aligns individual dimensions with fitness-relevant phenotypes such as antibody escape, ACE2 binding and expression. (**C**) The framework supports a broad range of evolutionary forecasting tasks, including pre-pandemic prediction of high-risk mutations, identification and mechanistic interpretation of antigenic or functional risk sites, reconstruction and foreshadowing of evolutionary trends and generalization across antigenically and structurally diverse viral families.

Next, the multimodal evolutionary history is used as input for disentangled representation learning within a variational autoencoder (VAE) architecture (*28*), which provides a tractable approximation to the underlying evolutionary distribution. Conventional VAEs employ a fixed isotropic Gaussian prior, which imposes a symmetric and single-peaked latent structure, to learn a unimodal embedding, and this formulation has been widely used to approximate mutational fitness through likelihood-based scoring (*29, 30*). However, such a prior does not reflect the multipeaked and heterogeneous nature of viral evolutionary pressures. Fitness-relevant factors such as folding stability, receptor binding, expression and immune evasion occupy distinct and sometimes opposing regions of sequence space and collapsing them into a single unimodal prior can mix their effects, bias likelihood estimates and obscure biologically meaningful variation. To address this limitation, DERIVE replaces the fixed Gaussian prior with a learnable, factorized normalizing-flow (*31*) prior regularized by a total-correlation penalty. The flow prior consists of a sequence of invertible, smooth transformations that reshape the latent space to better match the posterior geometry implied by the multimodal evolutionary history (Fig. 1B). The resulting latent space is flexible, multimodal and anisotropic, allowing different biological factors to occupy distinct regions rather than being conflated. The total-correlation regularization further promotes disentanglement among latent dimensions or features, reducing coupling between antigenicity-linked variation and other evolutionary pressures such as stability or receptor affinity (Methods).

With this design, DERIVE captures a disentangled physicochemical–structural evolutionary distribution, enabling interpretable estimation of mutational effects, robust prediction under sparse-data conditions and improved generalization across divergent viral families. By modeling the multimodal structure of viral evolutionary constraints within a disentangled latent space, DERIVE supports a broad range of applications relevant to early antigenic assessment and public-health response, including (1) pre-pandemic prediction when experimental measurements are unavailable, (2) risk-site identification and mechanistic interpretation of antigenic determinants, (3) reconstruction of evolutionary trends and inference of antigenic pathways, and (4) cross-virus generalization to influenza, HIV, rabies, Chikungunya and other pathogens (Fig. 1C).

### DERIVE enables accurate prediction of SARS-CoV-2 risk variants from pre-pandemic modeling

To assess DERIVE under a realistic early-warning setting, we trained the model using only data available before January 2020, thereby emulating pre-pandemic constraints in which no SARS-CoV-2–specific assays or functional annotations existed (**Methods**). Using this pre-pandemic model, we calculated DERIVE scores for all 24,187 possible single–amino-acid substitutions in Spike and ranked them. We then compared this ranking with mutations reported in the GISAID database (*32*). Substitutions that subsequently appeared during the pandemic are strongly enriched toward the top of the DERIVE ranking, and this enrichment increases with observed frequency: mutations detected >10k or >100k times show the most pronounced deviation from random expectation, followed by those seen >10 or >100 times (*P* value < 2.16e-16 for all comparisons, Fig. 2A). Despite never accessing post-2020 data, DERIVE prospectively highlights many substitutions that later emerged at scale. High-scoring substitutions predominantly localize to S1, with particular enrichment in the N-terminal domain (NTD) and the receptor-binding domain (RBD), with relative depletion in S2 (Fig. 2B), mirroring the regions where pandemic-era mutations eventually accumulated.

**Fig. 2:**
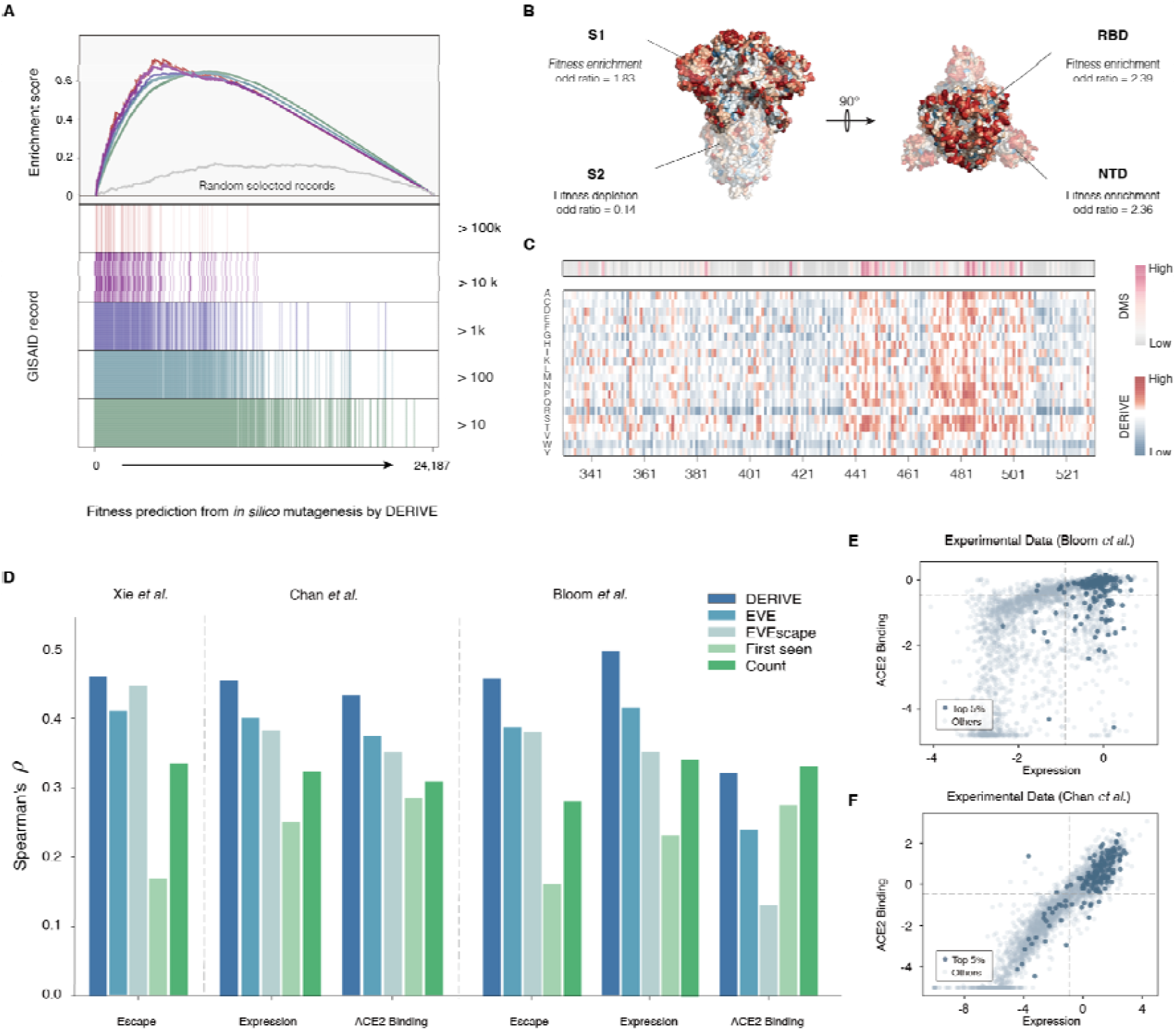
Performance of DERIVE in predicting SARS-CoV-2 risk variants from pre-pandemic modeling. (**A**) Enrichment of mutations that were later observed in GISAID, stratified by occurrence frequency. Fitness scores were obtained by *in silico* mutagenesis, where DERIVE scored all possible single amino-acid substitutions in Spike;the horizontal axis indexes mutations sorted by DERIVE fitness score and the vertical axis shows the enrichment score. Across all occurrence tiers (>100k, >10k, >1k, >100, >10), substitutions reported in GISAID are strongly and significantly enriched among high-scoring mutations inferred by DERIVE, whereas a random baseline shows no enrichment. (**B**) Regional enrichment measured by odds ratios, computed as the number of DERIVE high-scoring sites normalized by region length; values greater than 1 indicate enrichment and values less than 1 indicate depletion. (**C**) Concordance between DERIVE scores and experimental screening data in the receptor-binding motif (RBM). Heatmaps are arranged along the RBM sequence on the horizontal axis, with residue numbers indicating positions and rows corresponding to amino-acid substitutions; colour indicates the value at each site. The upper heatmap shows deep-mutational-scanning DMS measurements and the lower heatmap shows DERIVE scores at the same positions, highlighting aligned regions with high values. (**D**) Spearman correlations between model predictions and multiple DMS datasets for antibody escape, expression and ACE2 binding. The horizontal axis lists the phenotypes assayed in each study and the vertical axis shows Spearman’s ρ; bar colours distinguish DERIVE, EVE, EVEscape, first-seen date and GISAID count. DERIVE consistently achieves the highest correlations across these phenotypes. (**E** and **F**) DMS expression–ACE2-binding landscapes. Each point represents a single-mutant variant, with expression on the horizontal axis and ACE2 binding on the vertical axis. Variants ranked in the top 5% by DERIVE, shown in dark blue, cluster in the high-expression, high-binding quadrant compared with other variants shown in light grey.

Within the immunologically critical RBD, DERIVE prioritizes substitutions that satisfy well-quantified fitness constraints measured in independent deep-mutational scanning (DMS) assays (*33-43*). The top-ranked mutations from DERIVE cluster in the jointly high expression, high ACE2-binding region of the expression–ACE2 landscape (Fig. 2, E and F), and residues with high DERIVE scores coincide with experimentally measured antibody-escape effects from Bloom et al. assays (Fig. 2C). Both DERIVE and DMS analyses show concentrated signal in the receptor-binding motif (RBM) and relative depletion outside it. These consistencies indicate that DERIVE captures key biophysical principles governing RBD evolution without relying on SARS-CoV-2–specific training data.

Across multiple independent DMS datasets, DERIVE outperforms state-of-the-art baselines and heuristic surveillance indicators. It achieves the highest Spearman correlations with experimentally measured effects for antibody escape (Spearman’s ρ = 0.46 and 0.46 in Xie and Bloom et al.), RBD expression (Spearman’s ρ = 0.50 and 0.45 in Bloom and Chan), and ACE2 binding (Spearman’s ρ = 0.43 and 0.32 in Chan and Bloom) (Fig. 2D). These correlations substantially exceed those of EVE (*29*) (geometric mean 19%, median 17%) and EVEscape (*30*) (geometric mean 35%, median 21%), and clearly surpass simple predictors such as first-seen date or the GISAID report count. The stability of these gains across laboratories, phenotypes and assay designs suggests that DERIVE provides a more faithful ordering of single-mutation effects than existing models or surveillance statistics.

### Interpretable latent evolutionary determinants learned by DERIVE

To assess the interpretability afforded by disentanglement, we examined the latent features learned by DERIVE and compared them with multiple independently measured phenotypes relevant to viral fitness (Fig. 3A). Among the latent coordinates, the four dimensions with the largest variance showed stable correspondence to known biophysical determinants across multiple datasets (Supplementary Methods). The first latent feature correlated strongly with RBD expression, the second aligned with ACE2 binding, the third captured antibody-escape effects, and the fourth was negatively associated with ACE2 binding. Principal component analysis of the latent space further showed that mutations with high and low DERIVE scores occupy distinct regions, and that mutations annotated by expression, ACE2 affinity or antibody escape segregate into partially non-overlapping clusters (fig. S1). These observations indicate that DERIVE recovers separable, fitness-linked axes that reflect distinct and in some cases competing evolutionary pressures.

**Fig. 3:**
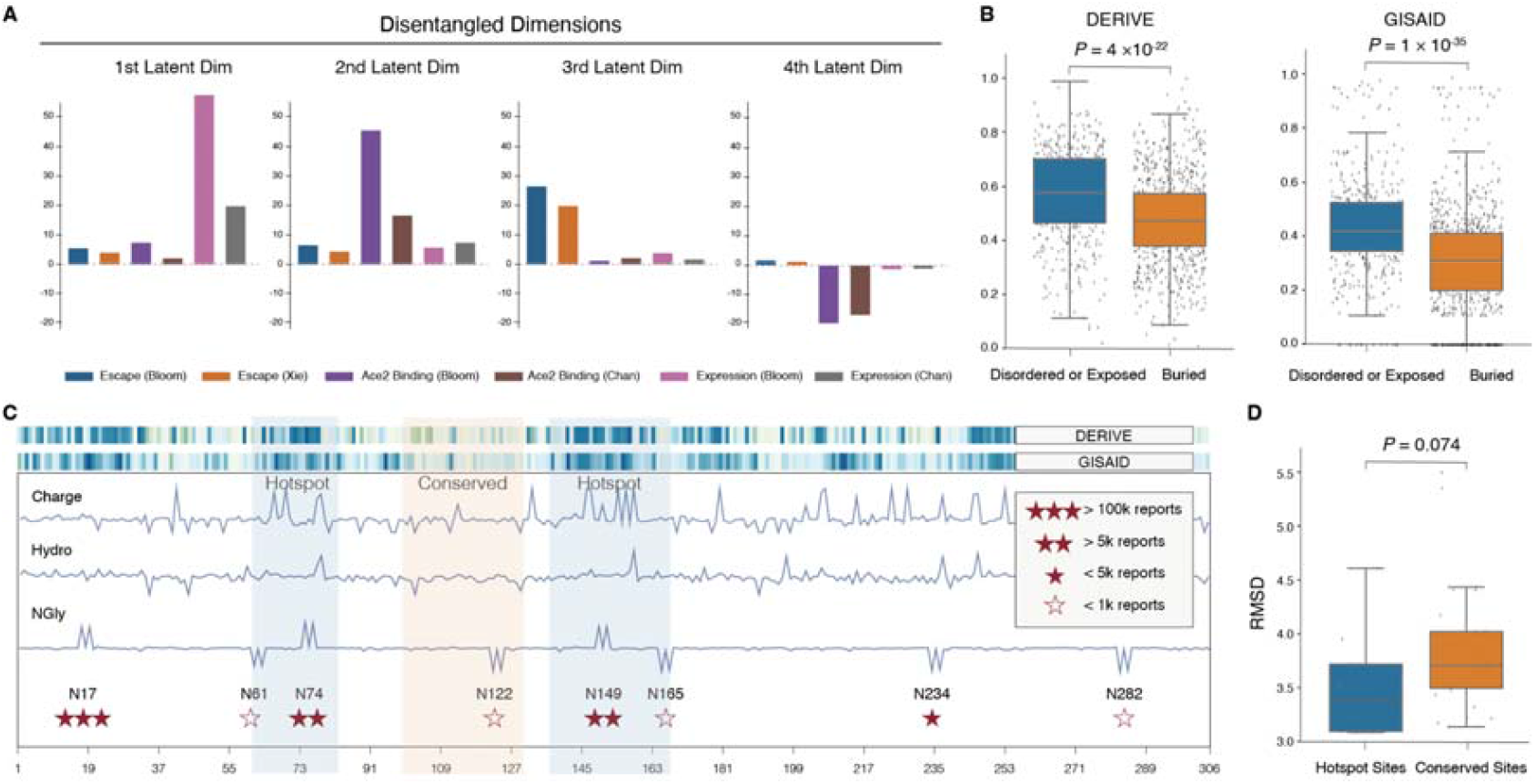
Interpretable disentangled factors and multimodal constraints learned by DERIVE. (**A**) Bar plots display the four latent dimensions with the highest variance. For each dimension, coloured bars indicate independent experimental phenotypes such as expression, ACE2 affinity and antibody escape measured in different datasets, and the vertical axis shows the LASSO regression coefficient linking that latent dimension to each phenotype, showing that these high-variance dimensions strongly correspond to the experimental measurements and indicating that DERIVE recovers separable axes of evolutionary variation linked to fitness. (**B**) Each point represents a Spike amino acid residue, grouped on the horizontal axis as disordered or solvent-exposed versus buried. The vertical axis shows mutational preference values, and the boxplots summarize the distributions of mutational preferences inferred by DERIVE on the left and the corresponding positional biases observed in GISAID on the right, showing that preferences inferred by DERIVE are enriched at disordered or exposed residues and mirror the biases seen in GISAID, consistent with accurate learning of spatial constraints on mutational tolerance. (**C**) Along the NTD sequence, with the horizontal axis giving residue index, the top two heat-map rows show the DERIVE score and the mutation signal from GISAID at each site, with darker colours indicating larger values. The three traces below show the contributions of charge, hydrophobicity and N-linked glycosylation to the DERIVE score on the vertical axis, expressed as relative contribution, with shaded bands marking mutation-sensitive hotspot regions and conserved regions, and asterisks marking sites repeatedly observed during the pandemic. Together, these multimodal signals delineate mutation-sensitive hotspots and conserved regions in the NTD, with glycosylation-associated features aligning with frequently observed sites. (**D**) Boxplots compare substitutions at hotspot sites and conserved sites highlighted in **C**, with the horizontal axis separating hotspot and conserved groups and the vertical axis giving backbone root-mean-square deviation (RMSD) values from molecular dynamics simulations. The simulations show that higher-scoring substitutions in hotspot regions identified by DERIVE have lower backbone RMSD than substitutions in conserved regions, consistent with greater structural stability.

We next evaluated how the physicochemical–structural evolutionary history contributes to these patterns. In the NTD, DERIVE identifies charge, hydrophobicity and N-linked glycosylation as influential features, and mutations highlighted by DERIVE align with pandemic hotspots (Fig. 3C). DERIVE correctly analyzes the influence of loss of N-linked glycosylation to virus evolution. Even though glycans often shield epitopes and promote immune evasion, DERIVE does not simply believe “loss glycan = worse”. For instance, loss of N17, N74, caused by substitutions such as T19I and T76I, is scored as beneficial by DERIVE and was later observed in BA.2 (Omicron) and its descendants, including recently spreading LP.8.1 and NB.1.8.1; and C.37 (Lambda) lineages, respectively. In contrast, DERIVE assigns deleterious effects to loss of glycans at N61, N122, N165, N234 and N282, consistent with experimental evidence showing reduced infectivity or diminished ACE2 binding when these sites are disrupted (*44-47*). When glycosylation, charge and hydrophobicity are analyzed jointly, the resulting signatures delineate a conserved region within the NTD that remains stable across Alpha, Beta and Omicron lineages and overlaps the footprints of broadly neutralizing NTD antibodies XG2v046 and XGv280 (*48*), indicating DERIVE’s potential utility for broad□spectrum antibody design.

We next asked whether the structural descriptors reflect biophysically plausible structural consequences and further enhance DERIVE’s ability to represent realistic mutational tendencies. Solvent exposure and intrinsic disorder modulate protein–protein interactions, as residues that are disordered or solvent-exposed are more likely to participate in immune recognition or receptor engagement (*49*). To emphasize these effects, DERIVE incorporates Gaussian-style data augmentation of GCN-derived features in disordered or solvent-exposed regions and employs a loss term that upweights these attributes. Comparison of DERIVE’s regional preferences with natural mutation frequencies observed in GISAID shows consistent enrichment in disordered and exposed sites (Fig. 3B), suggesting that the structural history is well captured in a qualitative manner.

In addition, we performed molecular dynamics (MD) simulations on single-amino-acid substitutions arising from triple-nucleotide changes, a mutation class too rare for reliable epidemiological validation. Using three MD replicas for each substitution in the folded NTD domain, substitutions with high DERIVE scores in hot regions (colored in Fig. 3C) exhibited lower backbone RMSD values than those with low scores in the reserved region, indicating greater structural stability (Fig. 3D). This difference indicates a relative advantage in structural stability for high-scoring mutations and reveals a broadly consistent trend. These analyses provide an independent, simulation-based validation of DERIVE’s ability to prioritize mutations that preserve structural integrity, supporting its use in early antigenic assessment and in guiding candidate selection for therapeutic development.

### Forecasting real-world evolutionary trajectories of SARS-CoV-2

A key application of DERIVE is supporting pandemic management under dynamic, real□world conditions. To evaluate DERIVE in conditions that approximate real-world viral surveillance, we generated monthly predictions in a time-updated manner, using training sets expanded and reweighted to include all data available before each prediction point (Methods). Longitudinal tracking shows that, with natural fluctuations, DERIVE increases the priorities of relatively new mutations that subsequently appeared in GISAID within the following one-month interval (Fig. 4A). This improvement was consistent across ranking thresholds: the top 20% of predictions captured most subsequent mutations, the top 5% accounted for more than half of them, and the top 1%, although small in number, contained a disproportionate share of high-impact substitutions. The consistency of these patterns across thresholds suggests that DERIVE’s predictive performance arises from intrinsic evolutionary signals rather than threshold-specific artifacts.

**Fig. 4:**
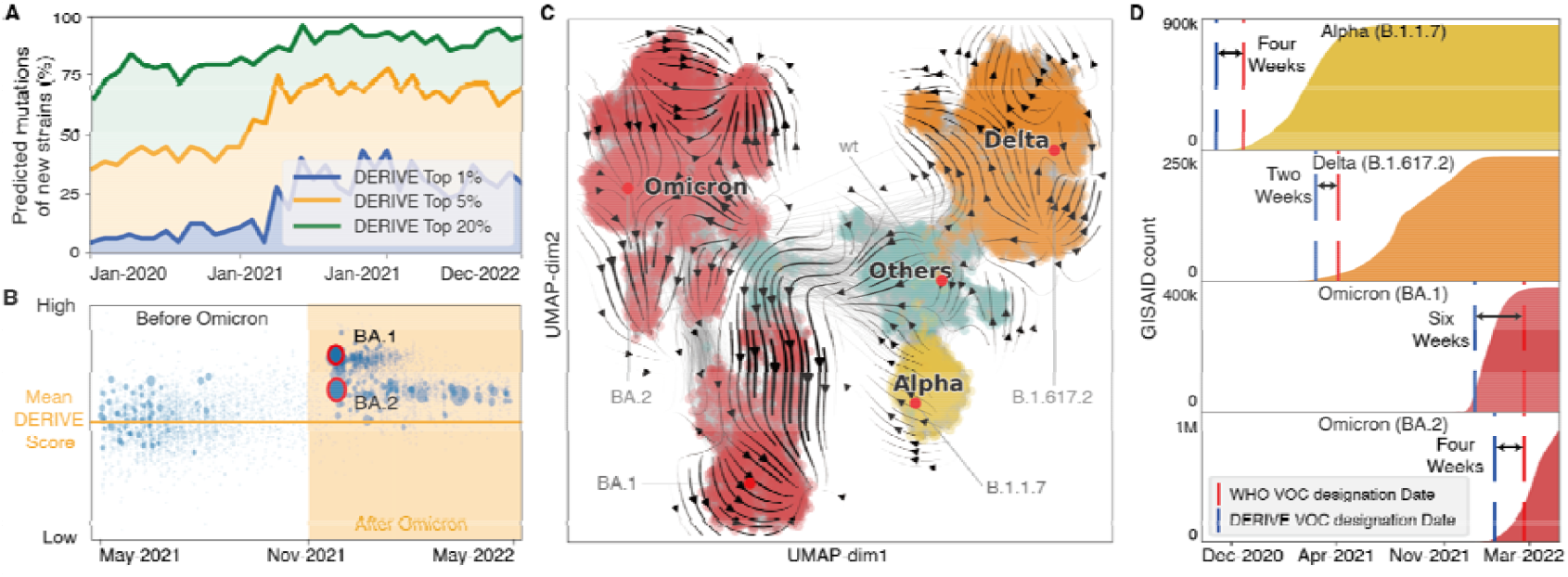
Prospective evolutionary forecasting and early variant detection for SARS-CoV-2. (**A**) The horizontal axis shows time in months and the vertical axis shows the proportion of mutations that newly appear in strains in the following month and are already contained in the current month’s high-scoring DERIVE subset. The three curves correspond to mutation subsets ranked in the top 20%, 5% and 1% of DERIVE scores in that month, based on monthly updated DERIVE predictions, showing that high-scoring subsets can anticipate a large number of new mutations and that enrichment increases at more stringent thresholds. (**B**) The horizontal axis shows sampling time and the vertical axis shows the strain-level DERIVE score, obtained by averaging mutation-level prediction scores across Spike substitutions. The vertical line separates the periods before and after the emergence of Omicron, and the BA.1 and BA.2 lineages are highlighted in the high-scoring region after Omicron, showing that scores increase after Omicron and that BA.1 and BA.2 lie above the pre-Omicron baseline. (**C**) Points are shown in a two-dimensional UMAP projection and coloured regions mark the Alpha, Delta, Omicron and other lineages. Black streamlines show evolutionary trajectories reconstructed by DERIVE, forming coherent and directed paths that coincide with major lineage transitions and highlighting the constrained nature of the adaptation routes. (**D**) Each row corresponds to one variant of concern: Alpha, Delta, Omicron BA.1 and Omicron BA.2. The horizontal axis shows time and the vertical axis shows the number of GISAID sequences for that variant. Coloured areas show how the counts for each variant change over time; red and blue vertical lines mark, respectively, the WHO designation date and the designation date from a biweekly early-warning procedure based on DERIVE predictions and lineage metadata, and black annotations indicate the interval between these dates, showing that this procedure can flag major variants several weeks before their WHO designation.

Because public-health actions are typically organized at the strain level (e.g., vaccines targeting, World Health Organization alerts), we next aggregated mutation-level predictions into a strain-level DERIVE score by averaging across all valid Spike substitutions in each lineage. Visualization from May 2021 to May 2022 (Fig. 4B) showed that the emergence of Omicron was accompanied by a marked elevation in strain-level scores, particularly for BA.1 and BA.2, relative to pre-Omicron lineages. This increase aligns with established evolutionary divergence and indicates that DERIVE is capable of prospectively flagging lineages with elevated evolutionary potential.

To examine broader evolutionary dynamics, we projected SARS-CoV-2 genome sequences into a reduced-dimensional evolutionary space to reconstruct evolutionary trajectories (Methods). Major lineages, including Alpha, Delta and Omicron, formed distinct and coherent trajectories that align with high-risk paths predicted by DERIVE (Fig. 4C). Directional flows indicated by black arrows show that viral evolution progresses along constrained adaptive corridors rather than through random diffusions, providing macro-scale support for the interpretability of the DERIVE scoring landscape.

Next, we implemented an operational early-warning procedure that integrates DERIVE predictions with lineage-level metadata from GISAID to produce a list of candidate Variants of Concern (VOCs) on a biweekly schedule (Methods). Comparison with official World Health Organization (WHO) designations showed that DERIVE flagged major VOCs several weeks earlier: Alpha 4 weeks earlier, Delta 2 weeks earlier, Omicron BA.1 6 weeks earlier and Omicron BA.2 4 weeks earlier (Fig. 4D). All these analyses show that DERIVE not only recapitulates historical evolutionary trajectories but also provides actionable foresight into the emergence of high-impact variants, highlighting its potential as an early warning system for global public health surveillance and pandemic preparedness.

### Cross-virus generalization of DERIVE across SARS-CoV-2, influenza, HIV, rabies and chikungunya

To systematically evaluate DERIVE’s ability to generalize across antigenically and structurally distinct viruses, we assembled a panel of viral antigens that differ substantially in sequence composition, structural organization and host-receptor engagement. We first examined influenza A virus hemagglutinin (HA) and the HIV-1 Env glycoprotein, two surface-exposed antigens with marked structural and functional divergence. For both proteins, the full DERIVE architecture, which integrates the disentangled representation module and the physicochemical feature module, achieved the highest Spearman correlations between model-predicted escape scores and replication-related phenotypes measured by DMS (Fig. 5A). Removing either module substantially reduced agreement with DMS, indicating that both inductive biases contribute essential and complementary information. These findings demonstrate that DERIVE maintains predictive consistency even when applied to antigens sharing minimal similarity in sequence or structure.

**Fig. 5:**
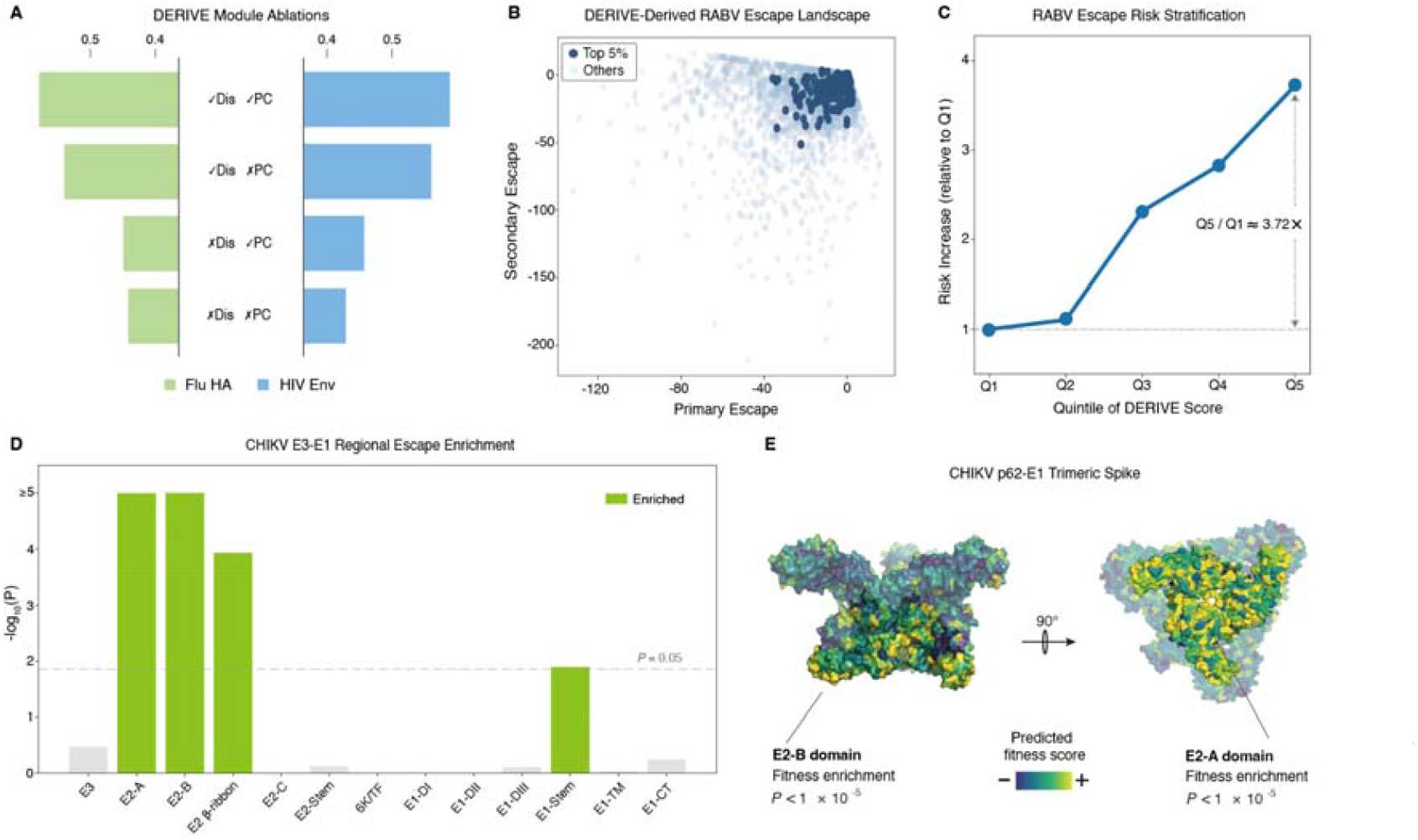
Cross-virus transferability of DERIVE in antigenic escape mapping. (**A**) Bar plots show ablation analyses of DERIVE performance for influenza A HA (left) and HIV-1 Env (right), comparing models with and without the disentanglement (Dis) and physicochemical (PC) modules; checkmarks and crosses indicate the presence or absence of each module. The vertical axis lists the model configurations and the horizontal axis shows Spearman correlation ρ between DERIVE-predicted escape scores and deep-mutational-scanning replication phenotypes. (**B**) Model-derived escape landscape for the rabies lyssavirus glycoprotein G. Each point corresponds to a single-amino-acid substitution, with the horizontal axis giving the primary escape axis obtained from DERIVE and the vertical axis giving an orthogonal secondary escape axis that summarizes residual variation; dark points denote the top 5% of variants by predicted escape score. (**C**) Risk stratification of CTB012 escape using DERIVE scores. Variants ranked by predicted escape score are divided into quintiles Q1–Q5 along the horizontal axis, and the vertical axis reports the relative risk of experimentally defined high-escape mutations in each quintile normalized to Q1. (**D**) Regional enrichment of high-scoring mutations on the chikungunya virus CHIKV p62–E1 spike, with the horizontal axis showing annotated structural domains and the vertical axis showing −log10(*P*) from enrichment tests; green bars indicate significantly enriched regions. (**E**) DERIVE escape scores are mapped onto the three-dimensional CHIKV p62–E1 trimeric structure PDB 6JO8, with surface colour indicating predicted escape potential and two views separated by a 90° rotation, highlighting dense clustering of high-escape mutations in the E2-A and E2-B domains and the connecting β-ribbon.

We next assessed the model on rabies lyssavirus glycoprotein, using a DMS dataset covering nearly all single–amino-acid substitutions. Model-derived escape scores were projected into a reduced-dimensional space aligned with the latent representation. A primary escape axis was constructed to maximize separation between the top-scoring variants (top 5%) and all remaining substitutions. High-scoring variants formed a well-defined cluster along this axis (Fig. 5B), indicating that DERIVE has learned a coherent escape gradient. Residual variation orthogonal to this axis was summarized by the first principal component in the residual space, revealing finer-scale organization among variants with similar escape potential. Together, these axes defined a two-dimensional escape landscape that captures the global topology of mutational effects predicted by the model.

To test the practical relevance of the escape scores for antibody-escape phenotypes, we performed a risk-stratification analysis using DMS data for the monoclonal antibody CTB012 (Fig. 5C). Variants were ranked by DERIVE scores and partitioned into five equal-sized score quintiles (Q1–Q5, from lowest to highest). The proportion of experimentally defined high-escape mutations increased monotonically from Q1 to Q5, reaching a 3.7-fold enrichment in the highest-scoring quintile Q5 relative to the lowest-scoring quintile Q1. This stratification demonstrates that the model-derived escape score provides a meaningful quantitative metric for prioritizing mutations with elevated immune-escape potential.

We then applied DERIVE to chikungunya virus (CHIKV), which is currently locally circulating, and analyzed escape patterns on the p62–E1 trimeric spike. High-scoring mutations were strongly concentrated in discrete regions of the E2 subunit (Fig. 5D). Domain-level analysis showed significant enrichment in the E2-A and E2-B domains (*P* < 10□□) and in the β-ribbon linking them (approximately *P* ≈ 10 □□). Other regions, including most of E1, showed values near background except for moderate enrichment in the E1-stem domain. These enriched regions correspond closely to known antigenic surfaces identified through epitope mapping of CHIKV and related alphaviruses (*50*). To visualize the spatial organization of these signals, we mapped DERIVE escape scores onto the three-dimensional p62–E1 structure (PDB ID: 6JO8), revealing dense clustering of high-scoring mutations on E2-A, E2-B and the connecting β-ribbon (Fig. 5E). These findings confirm that the model accurately highlights surface-exposed regions that constitute major neutralizing epitopes.

Taken together, these analyses show that DERIVE generalizes robustly across viral species, antigen structures and experimental modalities. The model maintains consistent predictive performance across SARS-CoV-2, influenza A, HIV-1, rabies lyssavirus and CHIKV, and its outputs are concordant with experimental observations at multiple scales, from single-mutant effects in DMS assays to domain-level and structure-level antigenic organization. The resulting escape score is therefore both interpretable and broadly applicable, providing a generalizable metric for immune-escape surveillance, vaccine immunogen design and the screening and optimization of therapeutic antibodies.

## DISCUSSION

In this study, we introduce DERIVE, a unified framework that integrates multimodal evolutionary information into a disentangled latent representation, enabling direct modeling of antigenic and functional determinants across diverse viral systems. By separating antigenicity-linked signals from other evolutionary pressures, DERIVE provides a structured view of the mutational landscape that generalizes across species, experimental modalities and biological scales. Retrospective analyses of SARS-CoV-2 show that DERIVE can identify high-risk mutations and lineages prior to their population-level expansion. Furthermore, evaluations across additional virus families demonstrate that the learned representation is transferable to pathogens with distinct sequence compositions, structural architectures and immune environments.

Beyond predictive accuracy, interpretability is essential for actionable early-stage outbreak use. DERIVE’s disentangled latent space aligns individual axes with biological pressures such as expression, receptor affinity and antibody escape, revealing mechanistic structure without phenotype-specific supervision. The multimodal evolutionary history further allows individual modalities to be examined independently; for example, glycosylation signatures correctly anticipate recurrent losses such as N17 in recently circulating SARS-CoV-2 variants. These observations illustrate how expanding evolutionary modalities can uncover determinants of adaptive change and provide interpretable insight into evolutionary pathways.

A practical forecasting system must also accommodate rapidly updating surveillance sequences, experimental measurements and contextual metadata. DERIVE demonstrates strong real-time performance even when trained solely on GISAID sequences and can incorporate new evidence as it emerges. Integration of virological assays, epidemiological indicators and expert assessments offers a natural route toward adaptive refinement, and strategies such as reinforcement learning or active learning could enable DERIVE to update its evolutionary representation under distributional shift.

In addition, micro-level evolutionary inference from DERIVE can also inform macro-level public-health decisions. Insights learned at the mutation level can serve as a foundational component for decision systems operating in real-world outbreak settings. As large language models and multi-agent frameworks become increasingly capable of synthesizing evidence, interacting with human experts and coordinating complex tasks, integrating DERIVE as one module within such systems could enhance strategic planning. In particular, an agent ecosystem equipped with DERIVE’s evolutionary predictions could support early variant triage, optimize vaccine strain selection and enable more adaptive and data-driven policy responses during rapidly evolving epidemics.

In sum, DERIVE offers a generalizable and efficient approach for forecasting viral evolution by unifying multimodal evolutionary history with disentangled representation learning. Its consistent performance across divergent viral families and diverse functional measurements underscores broad applicability.

## METHODS

### Overview of the DERIVE framework

DERIVE is designed to deliver interpretable and transferable predictions of viral protein antigenic evolution under pre-pandemic, limited-data conditions. The architecture comprises two components: evolution history generation and disentangled latent modeling, that together form a unified pipeline from sequence, physicochemical and topological history to fitness estimation.

Specifically, we denote by ***x***_**0**_ the complete amino acid sequence of the target viral protein. For COVID-19, the target is the wild-type SARS-CoV-2 spike protein, which mediates recognition and entry into human cells and therefore governs viral transmissibility and pathogenicity. The ***s***-th amino acid is denoted 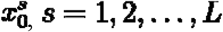, where ***L* = *l*(*x***_**0**_ ) is the length of the target sequence.

Through a two-stage sampling procedure, we obtain a discrete distribution ***p***^**∗**^ **(*x*)** from sequence-level homology search targeting ***x***_**0**_. Then we sample training dataset ***X***_***h***_ **= {*x***_***h***,**1**_, ***x***_***h***,**2**_,**…**, ***x***_***h***,***B***_**}** from the discrete distribution ***p***^**∗**^ **(*x*)** at each step ***h***, where ***B*** is the batch size. By augmenting and enhancing the data with physicochemical and topological information, we obtain the input dataset 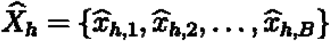, which will flow into the neural networks for training and analysis. For simplicity, we will omit the hat notation 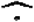 in the subsequent discussion of neural networks when no misunderstanding is likely to arise (i.e., use ***x*** instead of 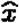, and ***X***_***h***_ instead of 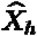). In short, evolution history generation can be summarized as the following:

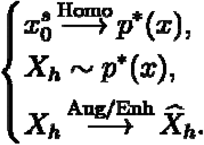

For each ***x*** in ***X***_***h***_, it passes through the encoder, which outputs ***μ***_***θ***_ **(*x*)** and ***σ***_***θ***_ **(*x*)**, where ***θ*** denotes all parameters of the encoder and decoder. Using the reparameterization trick proposed by Kingma and Welling^28^, we generate ***z* = *μ***_***θ***_ **(*x*) + *σ***_***θ***_ **(*x*) ⊙ *ϵ***, where ***ϵ*** is sampled from ***𝒩*(0,*I*)** and **⊙** is an element-wise product. After training, we have obtained the approximated continuous distribution ***p***_**θ**_ **(*x*)** for the discrete distribution ***p*\*(*x*)**. For any strain, the estimated fitness **Fitness(*x*)** is obtained via numerical estimation to ***p*_θ_(*x*)**, and it is proportional to **log *p*_*θ*_(*x*)**. The process from evolution history to fitness estimation can be written as:

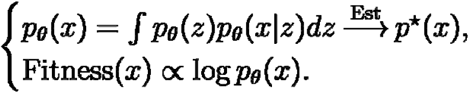

### Discrete formulation of evolutionary distribution from history

We train strictly on historical data to emulate pre-pandemic conditions. We aggregate four primary sources: UniRef100, UniRef90, MGnify, and BFD/MGnify. UniRef entries are restricted to those deposited before 2020-01-01, and we use the May 2019 release of MGnify (BFD/MGnify is constructed from this release). We then run jackhmmer-based multiple-sequence alignment for each target protein across these databases to retrieve homologous or functionally similar sequences, enforcing alignment quality via coverage (LCov) and the number of effective sequences (Neff).

We then construct the training set using a two-stage weighted sampling scheme. First, we perform probabilistic sampling at the database level. Second, we apply similarity-aware reweighting to adjust sample probabilities from each database and reduce oversampling bias, yielding a “balanced” evolutionary discrete distribution ***p*\*(*x*)**.

Specifically, let 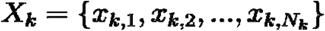 denote the aligned sequences from the k-th dataset (***k* = 1**, UniRef100; ***k* = 2**, UniRef90, ***k* = 3**, MGnify; ***k* = 4**, BFD/MGnify), where ***x***_***k,i***_ is i-th sequence from k-th dataset, and ***N***_***k***_ is the number of sequences of ***X***_***k***_. Then, the evolutionary discrete distribution ***p*\*(*x*)** is obtained by:

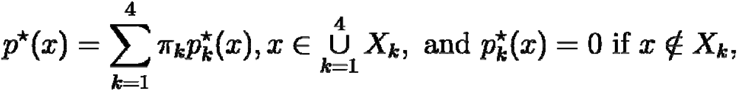

where ***π***_***k***_ is the weight for k-th dataset 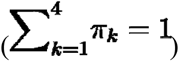, and 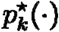 is the similarity □ aware rebalanced discrete distribution for ***X***_***k***_:

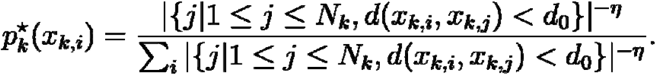

Here, ***d* (·,·)** is a function which makes **{*x***_***k***_, ***d* (·,·)}** a metric space, measuring diversity between aligned sequences, |***A***| denotes the cardinality of a finite set ***A***,|**{*j***| **1 ≤ *j* ≤*N***_***k***_, ***d*(*x***_***k,i***_, ***x***_***k,j***_**)< *d***_**0**_**}**| represents the local redundancy of ***x***_***k,i***_, and ***η*** is balancing strength (penalizing local redundancy), ranging from 0 to 1. Intuitively, ***η* = 0** recovers uniform sampling (no additional balancing) ***η* = 1** and works similar to cluster rebalancing method (firstly perform clustering, secondly sample uniformly across clusters, finally sample uniformly within the selected cluster).

Herein, we set ***d* (·,·)** be Hamming distance for simplicity 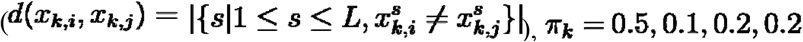 for ***k* = 1,2,34**, for, respectively, and choice of ***d***_**0**_ and ***η*** are provided in Ensemble learning in Methods.

### Multimodal Evolutionary Feature Construction

The previously derived MSA-based evolutionary distribution ***p*\*(*x*)** primarily provides a “sequence history,” capturing statistical patterns such as motifs that reflect amino acid types and covariation. However, protein–protein interactions, receptor engagement, and immune recognition are also governed by physicochemical and structural constraints beyond sequence—for example, hydrophobic cores are typically enriched in hydrophobic aliphatic residues; charged networks obey spatial compatibility and screening; and surface accessibility together with disorder modulates epitope exposure and conformational flexibility. Therefore, on top of the sequence history, we constructed a multimodal history that includes physicochemical attributes such as charge and hydrophobicity, coarse-grained force-field– derived statistics, as well as an indicator of N-linked glycosylation. This enables the model to learn not only which amino acids tend to co-occur, but also how those amino acids can physically coexist and contribute to function.

Beyond sequence-level context, protein interactions emerge from local structural neighborhoods. We model the reference protein’s 3D structure as a residue-level graph with two edge types—sequential and spatial. Given that short-range effects dominate residue interactions, we employ graph convolutional networks (GCNs) to aggregate neighborhood information, effectively mirroring this biophysical principle. In our setup, a lightweight GCN serves as a structural filter that fuses the physicochemical and glycosylation features over the graph, capturing the local milieu that constrains binding and recognition.

To mitigate potential data sparsity and variable MSA quality in future, as-yet-unknown viruses, we employ a biologically principled data enhancement scheme. Gaussian-style enhancement forces the model to focus on residues that are solvent-exposed or intrinsically disordered, which disproportionately mediate receptor binding, immune visibility, and conformational adaptability. Exposure is quantified by relative total solvent□accessible surface area (SASA) and negative weighted contact number (nWCN), while disorder is inferred from missing coordinates or sequence-based predictors. Enhancement strength increases monotonically with exposure/disorder, amplifying salient signals without disrupting overall structural coherence. (see Supplementary Methods for details)

### Disentangled factorized normalizing-flow bayesian variational autoencoder

We model the fitness landscape using a Bayesian variational autoencoder (VAE)^28^, simultaneously imposing multiple constraints: accurate sequence reconstruction, a normalizing-flow-based prior on the latent variables, explicit disentanglement of latent factors, and a mild prior on the network parameters.

Specifically, given discrete distribution ***q*\*(*x*)** representing multimodal evolution history, we optimize:

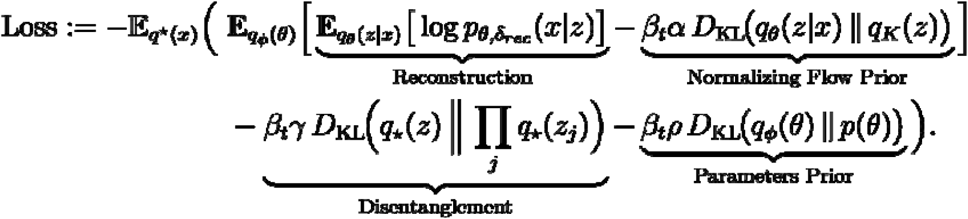

The terms in parentheses correspond to the reconstruction term, the normalizing-flow prior, the disentanglement regularizer, and the parameter prior, respectively. Here, ***θ*** denotes the encoder–decoder parameters. They are sampled from the parameter posterior ***q***_***ϕ***_**(*θ*), *ϕ*** with optimized by backpropagation; ***θ*** itself is not directly updated. Given ***θ***, the encoder induces the posterior ***q***_***θ***_ **(*z***|***x*)**. The distribution ***q***_***K***_**(*z*)** is a factorized normalizing flow composed of K invertible, smooth mappings, providing a flexible prior over latent variables that alleviates posterior mismatch. Disentanglement is encouraged by penalizing the total correlation between the aggregated posterior 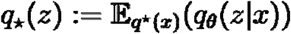 and the product of its marginals. The reconstruction term uses a position-weighted likelihood 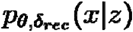 to emphasize functionally salient residues. The scalars ***β***_***t***_, ***α, γ***, and ***ρ*** control the weight of the flow prior KL, the disentanglement term, and the Bayesian parameter prior, balancing model expressivity, identifiability, and regularization under limited data.

#### Reconstruction Term

Guided by the observation that interactions with proteins such as antibodies or ACE2 are mediated by surface-exposed rather than buried residues, we use a position-weighted likelihood to emphasize such sites. Specifically, the decoder likelihood is Gaussian,

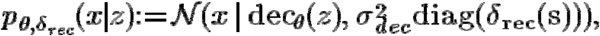

where ***δ***_***rec***_ **(*s*)** is a position weight for residue ***s*** that combines its disorder score ***D*(*s*)** and its degree of surface exposure, integrating the negatively weighted contact number (nWCN) and the relative total solvent-accessible surface area (SASA). The scalar 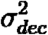 controls the overall strength of the reconstruction term.

#### Normalizing Flow Prior Term

We replace the vanilla VAE’s fixed Gaussian prior with a learnable, factorized normalizing-flow prior^31^.The flow prior is a sequence of invertible, smooth transformations that reshape the latent space to better match the posterior.

Specifically, we consider an invertible map ***f*** with inverse ***g*** that transforms a random variable ***z*** with density ***q*(*z*)**. The transformed density admits a closed form via the change-of-variables rule: 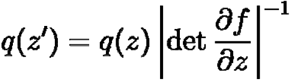. By factorizing the transformation into a composition of simple, tractable maps, ***f***_***K***_**○ *f***_***K*−1**_**○…○*f***_**1**_, we obtain ***z***_***K***_ **= *f***_***K***_**○ *f***_***K*−1**_**○…○*f***_**1**_**(*z*)** with density ***q***_***K***_ (**·**)and a concise log-density expression:

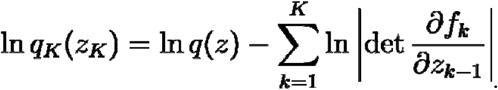

This construction replace the vanilla VAE’s normal prior with the flexible ***q***_***K***_ while keeping computation tractable, because each Jacobian determinant is easy to compute (e.g., triangular or low-rank structure).

#### Disentanglement Term

We introduce total correlation as a disentanglement regularizer to promote statistically independent latent factors. Specifically, we define total correlation ***TC* (*z*)** as

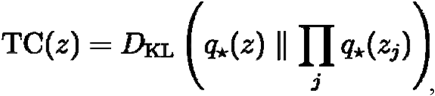

which penalizes dependencies among coordinates of the aggregate posterior ***q*_⋆_ (*z*)** and drives the model toward factorized representations. Note that **TC (*z*) = 0** if and only if all dimensions of the aggregate posterior are mutually independent, yielding a clearer and more separated latent structure. In practice, to obtain a non-negative, tractable estimator, we approximate total correlation by its lower bound, following Chen’s work in 2018 (*51*).

#### Parameters Prior Term

To mitigate overfitting, we treat network weights as random variables with a learnable parameter prior, optimized by backpropagation.

Specifically, we define the prior over parameters as 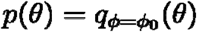, where ***ϕ***_**0**_ is a fixed constant setting for ***ϕ***. This makes ***p*(*θ*)** interpretable as a reference variational distribution over parameters. For tractability, we assume independence across dimensions, yielding ***D***_***KL***_**(*q***_**ϕ**_**(*θ*)**|| ***p*(*θ*)) ∑*D***_***KL***_**(*q***_**ϕ**_**(*θ***_***i***_)|| ***p*(*θ***_***i***_)).We further decompose 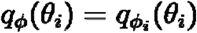, enabling position-specific regularization that adapts to each parameter’s role in the network.

If a parameter ***θ***_***i***_ lies in the encoder, or in the decoder but is not a bias term, we set 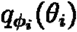 to be independent of ***ϕ***_***i***_. This choice implies ***D***_***KL***_**(*q***_**ϕ**_**(*θ***_***i***_) | | ***p*(*θ***_***i***_**)) = 0** for these parameters, so they receive no additional Bayesian regularization. This preserves the expressive capacity of feature extraction and weight-mediated decoding, while concentrating explicit prior control on parameters that set output baselines.

For decoder bias parameters, we introduce a tunable Gaussian variational family 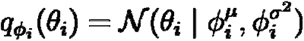, where 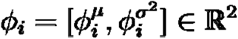. This grants decoder biases controlled freedom to adjust output offsets while being softly anchored by ***p*(*θ*)**. The same prior is shared across all decoder bias indices to maintain consistency.

### Ensemble learning

For small-scale unsupervised learning, performance can depend strikingly on the choice of random seeds, which controls training data generation (e.g. data enhancement), mini-batch sampling, weight initialization, hyperparameter selection and numerical approximation to fitness-related integral. This effect is especially pronounced when feature dimensionality is high and the sample size is limited. A systematic approach to mitigation is ensemble learning, which combines the results of multiple individual predictors, improving robustness, generalizability, and accuracy (*52, 53*). In practice, we found that using an ensemble of only ten networks sampled along the algorithm below, we achieved a clear improvement, across different tasks (see Supplementary Methods for details).

### Evolutionary distribution rebalancing with temporal awareness

When generating monthly, time-updated predictions, the key to success is the sophisticated construction of the training set: how we fold all data available before the prediction time into the evolving discrete distribution. We upgrade the prior similarity-aware reweighting (see Discrete formulation of evolutionary distribution from history) to a temporal-similarity-aware scheme that assigns higher weights to more recent variants. Intuitively, recent samples better anticipate the upcoming month, so contemporary epidemic strains should be learned preferentially.

Specifically, the epidemic-included evolutionary discrete distribution prior to time ***t*** is defined by:

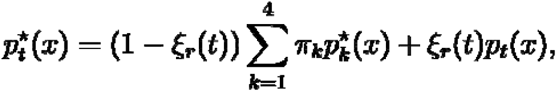

where 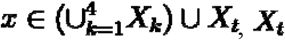 is a multiset with all spike amino acid sequences from the GISAID database submitted prior to time ***t, ξ***_***r***_ **(*t*)**, is a non-decreasing warm-up function, initialized at ***ξ***_***r***_ **(0) = 0.01**, exponentially increased until it reaches ***ξ***_***r***_ **(*t***_**0**_**) = 0.5** at time ***t***_**0**_ (Jan-2022, 24 months), and kept constant thereafter, and ***p***_***t***_(·) is temporal-similarity□aware rebalanced discrete distribution for ***X***_***t***_:

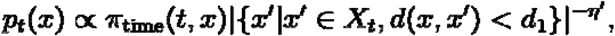

with the same neighborhood definition as 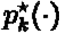. Here, ***d* (·,·)** be Hamming distance similar to 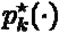, we let sequence similarity threshold ***d***_**1**_ **= 3**, balancing strength ***η***^***′***^ **= 0.25 × *η***, and temporal prior 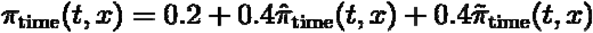, where 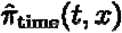 considers the first report time of x, 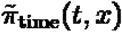 considers the final survival time of x, and both of them are exponentially increased from 0 to 1.

### Early detection of VOCs

We implemented an early-warning pipeline that produces a candidate VOC list every two weeks. At each time, we rank pangolin lineages by a composite score that rewards simultaneous prevalence, rapid expansion, and functional risk. To improve robustness, a lineage is included in the candidate VOC list only if it remains in the top five on every day over the 14-day interval ending at the cut-off. Each released candidate VOC list includes at most two candidates (often one), and the list may be empty.

Specifically, the composite score for lineage ***ℓ*** is ***S***_***ℓ***_ **(*t*)**, computed as a weighted geometric mean that integrates three components:

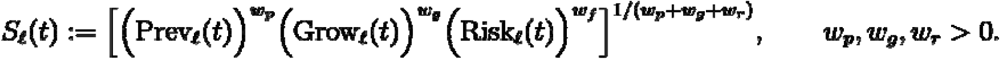

Here, contemporaneous prevalence **Prev**_***ℓ***_ **(*t*)** and growth score **Grow**_***ℓ***_ **(*t*)** were derived from cleaned GISAID data, and functional risk score **Risk**_***ℓ***_ **(*t*)** is estimated by DERIVE (see Supplementary Methods for details). Herein, we set ***w***_***p***_ **= 0.25, *w***_***g***_ **= *w***_***r***_ = **1**.

### Evolutionary trajectories reconstruction

To investigate the evolutionary dynamics of SARS-CoV-2 spike variants, we reconstructed evolutionary trajectories following the Evo-velocity framework introduced by Hie et al. (*54*), with adaptations to incorporate our DERIVE scores. We first collected all unique Spike amino acid sequences from the GISAID database submitted prior to May 1, 2023, that had been observed in the pandemic at least 50 times, ensuring that only high-confidence and epidemiologically relevant variants were included. Each sequence was embedded into a high-dimensional latent space using the pretrained protein language model ESM-1b^12^. To visualize the global structure of the sequence landscape, we applied Uniform Manifold Approximation and Projection (UMAP) to reduce the ESM-1b embedding space to two dimensions. Following Hie et al. (*54*), we constructed a K-nearest neighbor (KNN) graph based on the UMAP-reduced embeddings, where each node represents a unique spike sequence and edges connect each node to its K most similar neighbors in the embedding space. To assign directionality to the edges, we leveraged the DERIVE score for each sequence. For each directed edge between two nodes, we assigned the direction from the lower-scoring sequence to the higher-scoring sequence. This assumes that higher DERIVE scores correspond to evolutionarily favorable or antigenically advantageous variants, and that evolutionary trajectories tend to ascend this scoring landscape. The resulting vector field describes a DERIVE-driven prediction of antigenic evolution over the course of the pandemic.

## Acknowledgments

We are grateful to the members of the Yu Laboratory and many colleagues for their valuable comments and suggestions.

## Author contributions

F.Y. and Z.Z. conceived the project and designed the algorithm. Z.Z. and Y.L. implemented the algorithm and contributed to model training and data analyses. S.Z., P.Z. and Y.L. contributed to data interpretation and technical discussions. Z.Z., Y.L., and F.Y. wrote the manuscript with input from all authors. F.Y. supervised and directed the study.

## Data and materials availability

The databases Uniref100/Uniref90 are publicly available at https://ftp.uniprot.org/pub/databases/uniprot/uniref/. The database BFD/MGnify is publicly available at https://wwwuser.gwdg.de/∼compbiol/colabfold/bfd_mgy_colabfold.tar.gz. The database MGnify is publicly available at https://ftp.ebi.ac.uk/pub/databases/metagenomics/peptide_database/2019_05/. All SARS-CoV-2 sequencing and metadata data are available through https://gisaid.org/. The DMS datasets of binding affinity, expression and antibody escape prediction tasks of SARS-CoV-2 are publicly available from previous studies (*33-43*). The DMS datasets of the mutational effect prediction tasks of influenza virus, HIV, rabies and Chikungunya are also publicly available from previous studies (*55-58*). The source code for DERIVE is available at https://github.com/Yu-Lab-Genomics/DERIVE.

