## Supplemental Info for "Disentangled multimodal evolutionary representations for cross-virus predictive modeling of antigenic change"

Science Advances
AAAS

Supplementary Materials for

### Disentangled multimodal evolutionary representations for cross-virus predictive modeling of antigenic change

Zehong Zhang et al.

**This PDF file includes:**

Supplementary Text
Figs. S1 to S5

**Supplementary Text**

**Multimodal Evolutionary Feature Construction**

We first augment the amino acid sequence with non-structural properties—including physicochemical attributes and modification annotations. We then incorporate structural context and further enhance the structural representation by exposure level.

Specifically, after the discrete distribution $p^*(x)$ obtained, we can proceed to the generation of the training set. At each step $h$, a training dataset $X_h = \{x_{h,1}, x_{h,2}, \ldots, x_{h,B}\}$ is sampled from $p^*(x)$. (Note that we have index reuse here: $x_{k,i}$ represents the i-th aligned sequences from the k-th dataset; while, $x_{h,i}$ represents the i-th sequence from the training set sampled at step $h$, which could come from any dataset $k$.) Once the generation of the training set is completed, we will obtain the enhanced training set

$$\widehat{X}_h = [F_S(X_h), F_P(X_h), F_G(X_h), GF_A(X_h)]_F.$$

Here, $[\cdot, \cdots, \cdot]_F$ denotes data augmentation along the feature dimension (For example, if feature dimension is the third dimension, $A \in \mathbb{R}^{B \times L \times a}$ and $B \in \mathbb{R}^{B \times L \times b}$, then $[A, B]_F \in \mathbb{R}^{B \times L \times a+b}$). $F_S(\cdot)$, $F_P(\cdot)$, $F_G(\cdot)$ and $G(\cdot)$ represent sequence encoding, physicochemical augmentation, glycosylation augmentation and structural enhancement, respectively.

*Sequence information encoding*. Each amino acid residue $x^s_{h,i}, \forall h, i, s$ is vectorized through one-hot encoding into 20-dimensional sparse representations: $F_S(x^s_{h,i}) \in \mathbb{R}^{20}$, $F_S(X_h) \in \mathbb{R}^{B \times L \times 20}$. One-hot encoding is a method of representing categorical data as binary vectors, where only one element in the vector is 1 and the rest are 0. Unlike language-model-based implicit embeddings, this explicit encoding provides a transparent starting point, making the following feature interpretation and multimodal fusion more tractable.

*Physicochemical augmentation*. Inspired by AlphaFold3's use of charge as a model input, we observed that although neural networks can implicitly learn such features, the explicit introduction of specific physicochemical details significantly improves prediction accuracy.

In this work, physicochemical attributes—including charge (at physiological pH), hydrophobicity (Eisenberg-Weiss scale), polarity, and side-chain volume—were normalized and then assigned to each amino acid residue: $F_P(x^s_{h,i}) \in \mathbb{R}^4$, $F_P(X_h) \in \mathbb{R}^{B \times L \times 4}$. Furthermore, ablation studies and cluster analysis in the AAindex Database suggest that alternative feature selection and combination strategies may be viable. For instance, when we substituted polarity and side-chain volume with attributes derived from coarse-grained force-field statistics, the model achieved comparable performance. However, replacing charge and hydrophobicity degraded results. In summary, while researchers can adapt this augmentation strategy, features representing charge and hydrophobicity are essential and must be retained.

Attributes derived from coarse-grained force-field statistics. Following the FINCHES framework introduced by Holehouse et al. (*27*), we start from a set of analytical functions defined by a coarse-grained force field and end with the attributes for each residue.

Specifically, the total inter-residue non-bonded energy in the Mpipi coarse-grained force field (*59*) is defined as $M(p, q, r)$, where p and q denote the identities of amino acids from the 20 standard amino acids, and r is the distance between them. Then, the averaged interaction $E$ can be defined as:

$$E(p, q) := \int_{\sigma_{pq}}^{3R_{pq}} M(p, q, r)dr,$$

where $\sigma_{pq}$ and $R_{pq} := 3\sigma_{pq}$ are the zeros points of the Wang-Frenkel function with amino acid pair p and q. Here, an increase in the value of $E(p, q)$ indicates a decrease in attraction and an increase in repulsion between amino acids p and q.

To preserve the pairwise relations encoded by $E$ (as a matrix: $E \in \mathbb{R}^{20 \times 20}$) from the Mpipi coarse-grained force field, while reducing dimensionality, we employed non-metric multidimensional scaling (MDS) (*60*). This method preserves the rank order of pairwise dissimilarities by minimizing the stress within the target low-dimensional space. The resulting embedding thus yields low-dimensional attributes derived from the underlying force-field statistics. Alternatively, principal component analysis (PCA) yielded performance comparable to MDS.

*Glycosylation augmentation.* Glycosylation modifications, given their critical roles in receptor binding, immune shielding, and epitope accessibility, are augmented into input features as binary indicators. N-glycosylation is characterized based on canonical motifs (N-X-S/T, X≠P), while O-glycosylation is annotated according to literature support (*61*) (or neural network predictions (*62*) as substitution when related literature is absent): $F_G(x^s_{h,i}) \in \{0,1\}^2$, $F_G(X_h) \in \{0,1\}^{B\times L\times 2}$. Importantly, our analysis suggests that N-glycosylation must be included due to its ease of modeling and critical contribution to prediction accuracy, while O-glycosylation can be omitted.

*Structural enhancement.* The operator $G(\cdot)$ for structural enhancement is defined as:

$$G(\cdot) := E_{G,nWCN} G_2 E_{G,SASA} G_1(\cdot).$$

$G_i, i \in \{1,2\}$ represents graph convolutional neural networks (GCN) operation we employed, consisting of a pretrained GCN layer, with layer normalization and Leaky ReLU activation. Given limited training data, we adopt a simplified GCN that functions like a basic image filter: it scans each mutation's neighborhood (analogous to a 3×3 kernel) to integrate information between the mutated site and its surrounding protein environment. Concretely, we build this GCN as follows. First, we acquire the 3D structure of the protein (e.g. for SARS-CoV-2 Spike, we use the earliest publicly released PDB structure, 6VXX). Then, we construct an edge-weighted graph: a pair $(G,\omega)$, where the graph $G=(V,E)$ comprises residues as vertices $V$ and residue–residue links as edges $E$, and $\omega: E \to \mathbb{R}$ is a weight function. For each vertex $v \in V$ (corresponding to $x^s_{h,i}$) in the graph, we attach a 26-dimensional, sequence-physicochemical-glycosylation feature vector: $v := F_A(x^s_{h,i}) := [F_S(x^s_{h,i}), F_P(x^s_{h,i}), F_G(x^s_{h,i})]_F \in \mathbb{R}^{26}$. For each edge $e \in E$ (corresponding to connection between $x^s_{h,i}$ and $x^k_{h,i}$), we define its weight as $\omega(e) := \omega_{seq}(e) + \omega_{struc}(e)$. Specifically, if the sequential distance $d_{seq}$ between $x^s_{h,i}$ and $x^k_{h,i}$ (absolute index difference) is below a predefined threshold $D_{seq}$: $d_{seq}(x^s_{h,i}, x^k_{h,i}) \le D_{seq}$, these two residues are considered sequentially connected, and we set $\omega_{seq}(e) \propto d_{seq}(x^s_{h,i}, x^k_{h,i})^{-1}$; otherwise $\omega_{seq}(e) = 0$. If the spatial distance $d_{struc}$ between $x^s_{h,i}$ and $x^k_{h,i}$ (Euclidean distance between geometric centers of all non-hydrogen atoms of the residues; for glycines we use Cα coordinate) is below a predefined threshold $D_{struc}$: $d_{struc}(x^s_{h,i}, x^k_{h,i}) \leq D_{struc}$, these two residues are considered spatially connected and we set $\omega_{struc}(e) \propto d_{struct}(x^s_{h,i}, x^k_{h,i})^{-1}$. Herein, we set $D_{seq} = 2$ and $D_{struct} = 10\text{Å}$.

$E_{G,str}$, $\text{str} \in \{\text{SASA}, \text{nWCN}\}$ represents data enhancement. To deal with the limited training data, we apply a Gaussian-style augmentation, focusing on the disordered or exposed region (Considering $E_{G,SASA}$ first):

$$E_{G,SASA}(G_1(X_h)^s_i) = G_1(X_h)^s_i + \delta_{SASA}(s) \odot \epsilon, \ \epsilon \sim \mathcal{N}(0, I).$$

Here, $\delta_{SASA}(s)$ is the augmentation weight at position $s$. The weight increases with the disorder degree or exposure level, according to

$$\delta_{SASA}(s) = \begin{cases} 0, & S_T(s) < S_{T0} \text{ and } D(s) \leq D_1, \\ \alpha S_T(s), & S_{T0} \leq S_T(s) < S_{T1} \text{ and } D(s) \leq D_1, \\ \alpha S_{T1}, & S_{T1} \leq S_T(s) \text{ or } D_1 \leq D(s), \end{cases}$$

where $S_T(s)$ equals to relative total solvent-accessible surface area (SASA) at position $s$, $D(s)$ represents its disorder score (explained in the following section), $S_{T0}$, $S_{T1}$ and $D_1$ are predefined threshold, and $\alpha$ is the enhancement coefficient. In the same manner, we define $E_{G,nWCN}$, where the only difference is that the exposure level is quantified by the negative weighted contact number (nWCN) instead of SASA.

**Disorder and solvent exposure measures**

*Disorder score.* As a baseline estimation approach, the disorder score can be inferred from corresponding experimental structures. While observational biases (e.g., experimental resolution) exist, missing residue coordinates in the atomic model are often interpreted as evidence of intrinsic disorder: highly disordered segments exhibit conformational flexibility, leading to weak or absent experimental density that cannot be resolved into a single conformation. Consequently, these regions remain absent from the final structural model. Specifically, the disorder score $D(s)$ is defined as:

$$D(s) = \begin{cases} 0, & d(s, D_{\text{abs}}) \geq d_0, \\ \dfrac{d_0 - d(s, D_{\text{abs}})}{2d_0} + \dfrac{d(s, D_{\text{pdb}})}{2d_0}, & d(s, D_{\text{pdb}}) \leq d_0 \text{ and } d(s, D_{\text{abs}}) \leq d_0, \\ 1, & d(s, D_{\text{pdb}}) \geq d_0, \end{cases}$$

where $D_{\text{abs}}$ and $D_{\text{pdb}}$ represent the sets of amino acids with absent and experimentally resolved coordinates with respect to a certain structure, respectively. Clearly, $D_{\text{abs}} \cup D_{\text{pdb}}$ contains all amino acids and $D_{\text{abs}} \cap D_{\text{pdb}} = \emptyset$. The function $d(s, D) = \min\{d(s, s')|s' \in D\}$ is the distance from a point $s$ to a finite set $D$. Alternatively, Metapredict (*63, 64*) can estimate the disorder score when only sequence data is available, without requiring structural information.

*Solvent-exposure measures.* The core quantitative metrics for exposure degree can be categorized into two classes: geometric exposure, which includes the solvent excluded surface (SES) (*65*), solvent-accessible surface area (SASA) (*66*), and their many variations, including the relative solvent accessibility (RSA) (*67*); and local environmental density, which includes contact number (CN) (*68*) and its weighted counterpart, negative weighted contact number (nWCN) (*69*). Because metrics within a class are highly correlated: take SES and SASA for example, their difference chiefly lies in whether the surface is defined by the probe's center or by the inner envelope of the probe sphere. We select two representatives, one from each class, to avoid redundancy and maximize complementary information. Herein, we use the pair of relative total SASA and nWCN.

Specifically, relative total SASA $S_T(s)$ was calculated with FreeSASA (*70*), while nWCN $S_W(s)$ is defined by:

$$S_W(s) := \sum_{k \neq s} d_{struc}(x^k, x^s)^{-w},$$

where $d_{struc}$ denotes the spatial distance, calculated as the Euclidean distance between the geometric centers of residues k and s. These centers are computed over all non-hydrogen atoms. A special case is applied to glycine residues, which lack side-chain heavy atoms; for these, the position of the Cα atom is used as the geometric center. The exponent $w$ is a weighting parameter that controls the influence of local environmental density. Herein, we set $w = 2$.

*Disorder score and solvent-exposure measures combined in the reconstruction term.* Our reconstruction term, which focuses on the disordered or solvent-exposed, relies on the following weight function $\delta_{rec}(s) := \lambda_1\delta_{SASA}(s) + \lambda_2\delta_{nWCN}(s) + \lambda_3$, where $\delta_{nWCN}(s)$ is defined similar to $\delta_{SASA}(s)$. Herein, we set $\lambda_1 = \lambda_3 = 0.2$ and $\lambda_2 = 0.6$.

**Data cleaning**

We collected SARS-CoV-2 spike (S) protein sequences submitted to the GISAID database up to 1 May 2023. Several curation steps were applied. First, to focus on the human immune response to the virus, we retained only sequences derived from human hosts. Because protein synthesis typically initiates with methionine, we removed all S protein sequences that did not begin with methionine. Second, any sequence containing more than ten consecutive amino-acid substitutions was treated as a likely artifact and excluded. Finally, entries with duplicate accession IDs were considered redundant and removed.

**Latent-space variance and disentanglement analysis**

To assess whether individual latent dimensions capture distinct biological phenotypes, we analyzed the variance structure of the 64-dimensional latent space learned by DERIVE and its relationship to independent experimental measurements. For each RBD variant, DERIVE outputs a 64-dimensional latent representation vector $dim(z) \in \mathbb{R}^{64}$ in the latent space. We first computed the variance of each latent coordinate across all variants and ranked the 64 dimensions by their variance, and then selected the four dimensions with the highest variance (referred to Fig. 3A) for further analysis.

**Assessing latent disentanglement via Lasso coefficients**

For each experimental phenotype (for example antibody escape, expression or ACE2 binding), we fitted a separate L1-regularised linear model (Lasso) (*71*) that represents the phenotype as a linear combination of the latent dimensions. Let the latent vector of variant $i$ be $\mathbf{z}_i = (z_{i1}, \ldots, z_{iD})$ and the corresponding experimental measurement be $y_i$. Here, $N$ denotes the number of variant samples and $D$ the dimensionality of the latent representation. The Lasso model predicts

$$\hat{y}_i = \beta_0 + \sum_{j=1}^{D} \beta_j z_{ij},$$

and the coefficients $\beta_j$ are estimated by minimising

$$\frac{1}{2N} \sum_{i=1}^{N} \left( y_i - \beta_0 - \sum_{j=1}^{D} \beta_j z_{ij} \right)^2 + \alpha \sum_{j=1}^{D} |\beta_j|$$

Before fitting, we applied z-score standardisation to each latent dimension $z_{\cdot j}$ and to each phenotype $y$ column-wise so that every variable has zero mean and unit variance, which makes the coefficients for a given latent dimension comparable across phenotypes. The regularisation strength $\alpha$ was selected by five-fold cross-validation, and the final model was then refitted on all samples to obtain the coefficients.

**Ensemble learning**

Ensemble learning structures DERIVE as a highly modular framework, helping mitigate the inevitable dependence on training data in AI modeling and easing hyperparameter selection in fully unsupervised settings (*72-74*). This relaxes the quality requirements on MAS, whose quantification is challenging especially when across divergent databases, and reduces the time spent on hyperparameter search. In sum, this ensemble makes DERIVE markedly more robust**.**

*Details of the ensemble construction.* This section provides the detailed construction of the ensemble described in the main text. Specifically, for each model, we sample the following key parameters: the random seed (left unspecified; auto-generated) and the GCN initialization (pretrained with probability $p = 0.5$, otherwise not pretrained). The hyperparameters $d_0$ (for sequence similarity threshold) and $\eta$ (for balancing strength) related to $p^*(x)$ generation, the latent-space dimension $\dim(z)$, the number of iterations $K$ for the factorized normalizing flow, and the regularization scales $\alpha$ (for normalizing flow prior term) and $\gamma$ (for disentanglement term) are drawn as:

$$\begin{cases} d_0 \sim 0.4\delta_{0.01L} + 0.5\mathbb{U}(0.01L, L), \\ \eta \sim 0.5\delta_1 + 0.5\mathbb{U}(0.25, 1), \\ \dim(z) \sim 0.5\delta_{32} + 0.5\mathbb{U}_{\{16,32,64,128\}}, \\ K \sim 0.3\delta_0 + 0.3\delta_8 + 0.5\mathbb{U}_{\{4,8,16,32\}}, \\ \alpha = \gamma \sim 0.4\delta_1 + 0.4\delta_2 + 0.2\mathbb{U}(1, 2), \end{cases}$$

where $\delta_x$ is the Dirac measure at $x$, $\mathbb{U}_A$ is the uniform distribution on a finite set $A$, and $\mathbb{U}(a, b)$ is the uniform distribution on the interval $(a, b)$.

With the parameters $\theta^{(i)}$ fixed, once training is completed, $i$-th ($i = 1, ..., M$) model produces an estimation $\log p_\theta^{(i)}$ to $\log p^\star$. These estimators are then normalized and aggregated:

$$\text{Fitness}(x) := \frac{1}{M}\sum_{i=1}^{M} \text{Normalizer}(\log p_\theta^{(i)}(x)),$$

where $M$ is the ensemble size. (larger $M$ generally improves performance. We set $M = 100$; if computation time is a limiting factor, $M = 10$ serves as a slightly less accurate trade-off.)

*Comparative evaluation of a 10-model ensemble versus single models.* We first trained 200 base learners as described in Methods (Ensemble learning). From these, we uniformly sampled 10,000 distinct ensembles, each aggregating 10 non-overlapping models. We then compared the performance of the 200 single models with the 10-model ensembles. For every model or ensemble, nine metrics were computed covering antibody escape, ACE2 binding, expression, and GISAID-based predictions, with two complementary visualizations (fig. S2). Histograms with kernel-density overlays depict the full performance distributions (single models in blue; ensembles in orange), while boxplots report medians and interquartile ranges pre- and post-ensembling for each task.

In sum, a simple 10-model ensemble consistently outperforms individual models across all evaluated tasks. The ensemble yields higher central performance and substantially reduced variance, leading to more reliable and generalizable predictions. Given the minimal methodological complexity and strong gains, we recommend adopting a modest ensemble as a default strategy when computational budget allows.

*Effect of Ensemble Size.* From the pool defined above (200 distinct models), we formed ensembles of size $M \in \{1, 2, 3, 4, 5, 10, 30, 100\}$. For each $M$, we repeatedly sampled

$M$ distinct members without replacement and aggregated their predictions by four aggregate scorers.

The enhancement of predictive accuracy is illustrated through the mean and median of the aggregate scores (fig. S3). Both metrics show a clear trend of monotonic improvement as $M$ grows. Specifically, the mean and median rise steeply from a single model ($M = 1$) up to small ensembles ($M = 10$), before transitioning into a phase of slower, sustained growth as the ensemble size increases further toward $M = 100$.

Regarding robustness gains, the analysis focuses on two key indicators: the minimum and the standard deviation of the aggregate scores (fig. S4). The elevation of the minimum score with increasing $M$ demonstrates improved reliability in the most challenging scenarios. Meanwhile, the near-linear reduction in standard deviation as a function of $\log(M)$, indicating markedly reduced variability as ensembles grow.

In conclusion, increasing ensemble size simultaneously improves accuracy and robustness. Most of the benefit is realized by moving from single models to small ensembles ($M \leq 10$); larger ensembles ($M \geq 30$) deliver smaller but consistent gains and tighter confidence. Practically, $M \in [10, 30]$ offers a favorable trade-off between performance and computational cost.

**Early detection of VOCs**

We built a biweekly early-warning pipeline that outputs a candidate VOC list. At each update, Pango lineages are ranked by a composite score combining prevalence, rapid growth, and functional risk. A lineage is nominated only if it stays within the daily top five throughout the preceding 14 days.

*Prevalence score* is defined by:

$$\mathrm{Prev}_{\ell}(t) = \begin{cases} \dfrac{N(W_t, \ell)}{N(W_t)}, & N(W_t) \geq N_0, \\ 0, & \text{otherwise}, \end{cases}$$

where $W_t$ is a 14-day window ending at time $t$, $N(W_t, \ell)$ is the number of reports assigned to lineage $\ell$ within $W_t$, and $N(W) := \sum_\ell N(W, \ell)$ is the total reports. We set the threshold $N_0 = 1000$.

*Growth score* is defined by:

$$\text{Grow}_\ell(t) = \begin{cases} 0, & \text{Prev}_\ell(t) \le P_0, \\ 2, & \text{Prev}_\ell(t) > P_0 \ \& \ \text{Prev}_\ell(t - T_0) \le P_1, \\ \dfrac{\text{Prev}_\ell(t)}{\text{Prev}_\ell(t - T_0)}, & \text{otherwise,} \end{cases}$$

where time gap $T_0 = 14$, threshold $P_0 = 10^{-3}$, and $P_1 = 10^{-4}$. Alternatively, a logistic regression model is commonly used to estimate relative growth advantage. To keep things simple and to accommodate heterogeneous sampling, we adopt this simplified approach.

*Risk score* is defined by:

$$\text{Risk}_\ell(t) = \max\left\{ \text{Fitness}(x) | o^{\text{Tukey}}_{W_t,\ell}(x) = 0, o^{\text{Z}}_{W_t,\ell}(x) = 0, \text{ and } x \in V(W_t, \ell) \right\},$$

where $V(W_t, \ell)$ is a set of $x$ reported in W_t and assigned to $\ell$, both $o^{\text{Tukey}}_{W_t,\ell}$ and $o^{\text{Z}}_{W_t,\ell}$ are outlier indicators by Tukey test and Z-score test, respectively:

$$o^{\text{Tukey}}_{W_t,\ell}(x) = \begin{cases} 1, & x > Q_{3,V(W_t,\ell)} + k(Q_{3,V(W_t,\ell)} - Q_{1,V(W_t,\ell)}), \\ -1, & x < Q_{1,V(W_t,\ell)} - k(Q_{3,V(W_t,\ell)} - Q_{1,V(W_t,\ell)}), \\ 0, & \text{otherwise,} \end{cases}$$

where $Q_{1,V(W_t,\ell)}$ and $Q_{3,V(W_t,\ell)}$ are the first and third quartiles of $V(W_t, \ell)$, respectively, and

$$o^{\text{Z}}_{W_t,\ell}(x) = \begin{cases} 1, & \dfrac{x - \mu_{V(W_t,\ell)}}{\sigma_{V(W_t,\ell)}} > Z_0, \\ -1, & \dfrac{x - \mu_{V(W_t,\ell)}}{\sigma_{V(W_t,\ell)}} < -Z_0, \\ 0, & \text{otherwise,} \end{cases}$$

where $\mu_{V(W_t,\ell)}$ and $\sigma_{V(W_t,\ell)}$ are the mean and standard deviation of $V(W_t,\ell)$, respectively. Herein, we set $k = 1.5$, $Z_0 = 2.5$.

**Molecular dynamics simulations**

*Initial structure preparation*. To evaluate the structural stability of SARS-CoV-2 mutations, we prepared initial structures based on PDB protein structures. We used the electron microscopy structure of the SARS-CoV-2 spike trimer (PDB ID: 6VXX), as the starting point. To reduce system complexity and computational cost, we focused on a structurally independent segment of the spike protein: residues 27-291 (NTD) of chain A. To generate the mutant structure (Take NTD-W104D for example), we basically retained the coordinates of backbone atoms (N, CA, C, O) of residue W104, introduced small perturbations to the backbone coordinates and explored alternative side-chain conformations. For each perturbed backbone, we used SCWRL4 (*75*) to generate the optimal side-chain rotamer of aspartate. This procedure yielded three distinct conformations of the NTD-W104D structures.

*System construction*. All systems were prepared using tleap from the AmberTools25 suite (*76*). Missing residues and hydrogen atoms were added, and protonation states were assigned under physiological pH. The ff19SB force field (*77*) was applied to the protein. Each system was solvated in an explicit OPC water box, extending 10 Å beyond the protein surface in all directions. Periodic boundary conditions were applied. To mimic physiological ionic strength, $Na^+$ and $Cl^-$ ions were added to neutralize the system and reach a final salt concentration of 150 mM NaCl.

*Energy minimization and equilibration*. We applied a multi-step equilibration protocol following standard practices based on Amber24 (*78*). Initially, energy minimization was performed under strong positional restraints on heavy atoms, first using constant volume (NVT) conditions without SHAKE constraints, and then with SHAKE applied to constraint bonds involving hydrogen atoms. Systems were gradually heated to 298 K using a Langevin thermostat, followed by pressure equilibration under NPT conditions using a Monte Carlo barostat. Restraint forces were progressively reduced over multiple equilibration phases. Once all restraints were removed, we performed 500 ns of production MD under NPT conditions to generate equilibrated conformations.

*Replica-exchange enhanced sampling with arbitrary degrees of freedom (REAF).* To overcome the limitations of conventional MD in sampling the conformational landscape—especially at physiological temperatures where systems are prone to becoming trapped in local energy minima—we applied Replica Exchange with Arbitrary Degrees of Freedom (REAF), a REST2-like (*79*) enhanced sampling technique implemented in AMBER. Compared to temperature replica exchange MD (T-REMD), REAF achieves efficient sampling with significantly fewer replicas by selectively scaling the solute–solvent interactions.

For each system, we ran REAF simulations with 10 replicas at the following temperatures: 300, 322, 345, 368, 394, 423, 455, 491, 529, and 572 K. Exchanges between neighboring replicas were attempted every 2 ps, resulting in an average acceptance rate of ~15% across all systems. The lowest-temperature replica (300 K) was used for subsequent calculations. All REAF simulations were performed under NPT conditions with the same thermostat and barostat settings as described above. The total simulation time for each replica was 500 ns, ensuring adequate sampling of conformational space for both wild-type and mutant complexes. RMSD values were calculated over the 200–500 ns portion of the trajectory, ensuring measurements reflect equilibrated dynamics.

**Ablation experiments**

We have evaluated the ablation studies of the disentangled representation module and the physicochemical feature module previously (Fig. 5A). In this part, the full DERIVE architecture includes the complete physicochemical attributes set (with CH: Charge and Hydrophobicity and PV: Polarity and side-chain Volume), the graph convolutional network with data augmentation (denoted as GCN), and Bayesian parameter priors (Bayesian VAE, denoted as θ prior). Initially, to test different combinations of physicochemical attributes, we replaced polarity and side-chain volume with counterparts derived from coarse-grained force-field statistics (+FF-PV), and subsequently replaced charge and hydrophobicity with attributes obtained from the same source (+FF-CH). Subsequently, we ablated the GCN module along with its associated data augmentation (-GCN), and subsequently removed the parameter priors (-θ prior). For architecture consistency, we tested 1,000 3-ensembled models of each ablation study.

Our analysis employed a consistent framework: nine complementary performance indicators were evaluated for every individual model, followed by an integrated visualization (fig. S5). Across all tasks, their performances are not strictly uniform, but there is a clear trend: DERIVE = +FF-PV ≥ +FF-CH ≥ -GCN ≥ -θ prior.

**Construction of model-aligned escape coordinates for the rabies glycoprotein**

To construct the model-aligned escape coordinates shown in Fig. 5B, we used deep mutational scanning data for the rabies lyssavirus glycoprotein. For each variant, we collected multiple antibody-related escape readouts and concatenated them into a feature vector $\mathbf{x}_i \in \mathbb{R}^D$. For each dimension, we applied a $\log(1+x)$ transformation and then performed robust standardization based on the median and median absolute deviation (MAD), yielding standardized feature vectors $\mathbf{z}_i$.

Based on the model-predicted escape scores, we used the 95th percentile as a threshold to define a high-scoring set (top 5%, denoted $y_i = 1$) and a remaining set ($y_i = 0$). Let $\boldsymbol{\mu}_1$ and $\boldsymbol{\mu}_0$ denote the sample means of the two classes in the standardized space, and let $\boldsymbol{\Sigma}$ be the empirical covariance matrix estimated over all variants. On this basis, we defined a ridge-regularized Fisher linear discriminant direction (*80*):

$$\mathbf{w} = \left(\boldsymbol{\Sigma} + \lambda \mathbf{I}\right)^{-1}\left(\boldsymbol{\mu}_1 - \boldsymbol{\mu}_0\right),$$

where $\lambda > 0$ is a regularization parameter. We then normalized this vector to unit length, $\hat{\mathbf{w}} = \frac{\mathbf{w}}{\|\mathbf{w}\|}$, and, if necessary, flipped its sign so that the mean projection of the high-scoring set along $\hat{\mathbf{w}}$ exceeded that of the remaining set. For each variant, the coordinate along the primary escape axis was defined as $s_{1,i} = \hat{\mathbf{w}}^\mathsf{T}\mathbf{z}_i$. To obtain a secondary axis orthogonal to this primary direction, we removed the component of $\mathbf{z}_i$ along $\hat{\mathbf{w}}$, yielding residual vectors $\mathbf{r}_i = \mathbf{z}_i - s_{1,i}\hat{\mathbf{w}}$. We then formed the covariance matrix over the residuals $\{\mathbf{r}_i\}$ and performed principal component analysis, taking the eigenvector corresponding to the largest eigenvalue as $\mathbf{v}$. The coordinate along the secondary escape axis was defined as $s_{2,i} = \mathbf{v}^\mathsf{T}\mathbf{r}_i$. In Fig. 5B, the scatter plot is drawn using $(s_{1,i}, s_{2,i})$ as the horizontal and vertical coordinates for all variants, where the primary axis maximizes the separation between model high-scoring variants and the remaining variants, and the secondary axis captures residual structured variation orthogonal to this escape gradient.

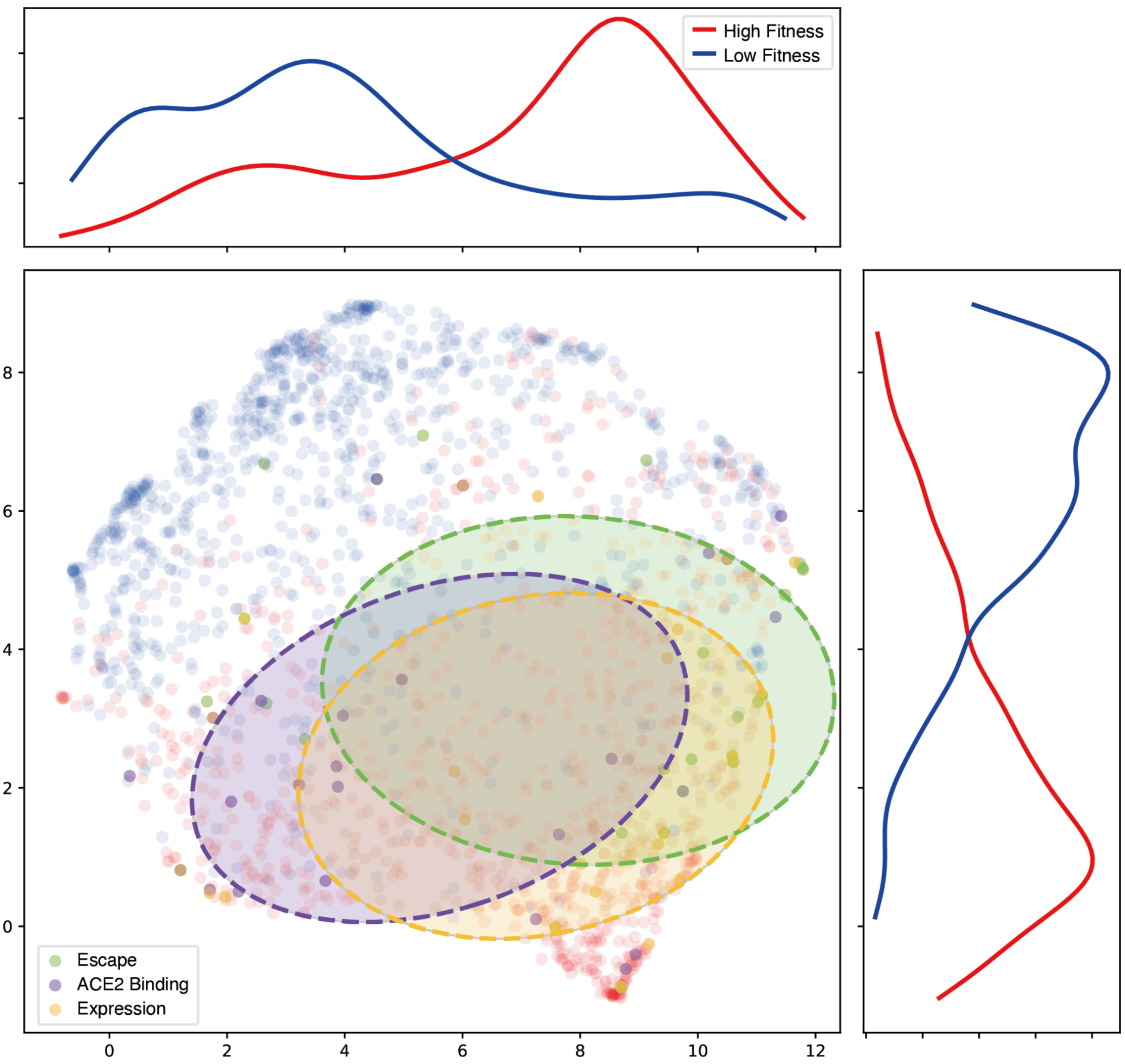


**Fig. S1. Dimensionality reduction visualization of latent space.** Central: UMAP embedding of learned latent representations. Mutations in the top 10% of the DERIVE score are shown in red; those in the bottom 10% are shown in blue. Mutations exhibiting a dominant effect for a single phenotype (escape, ACE2 binding, or expression) are additionally colored by that phenotype. Top/right: Kernel density estimates contrasting high- (red) and low-fitness (blue) mutations, showing clear separation between the two distributions.

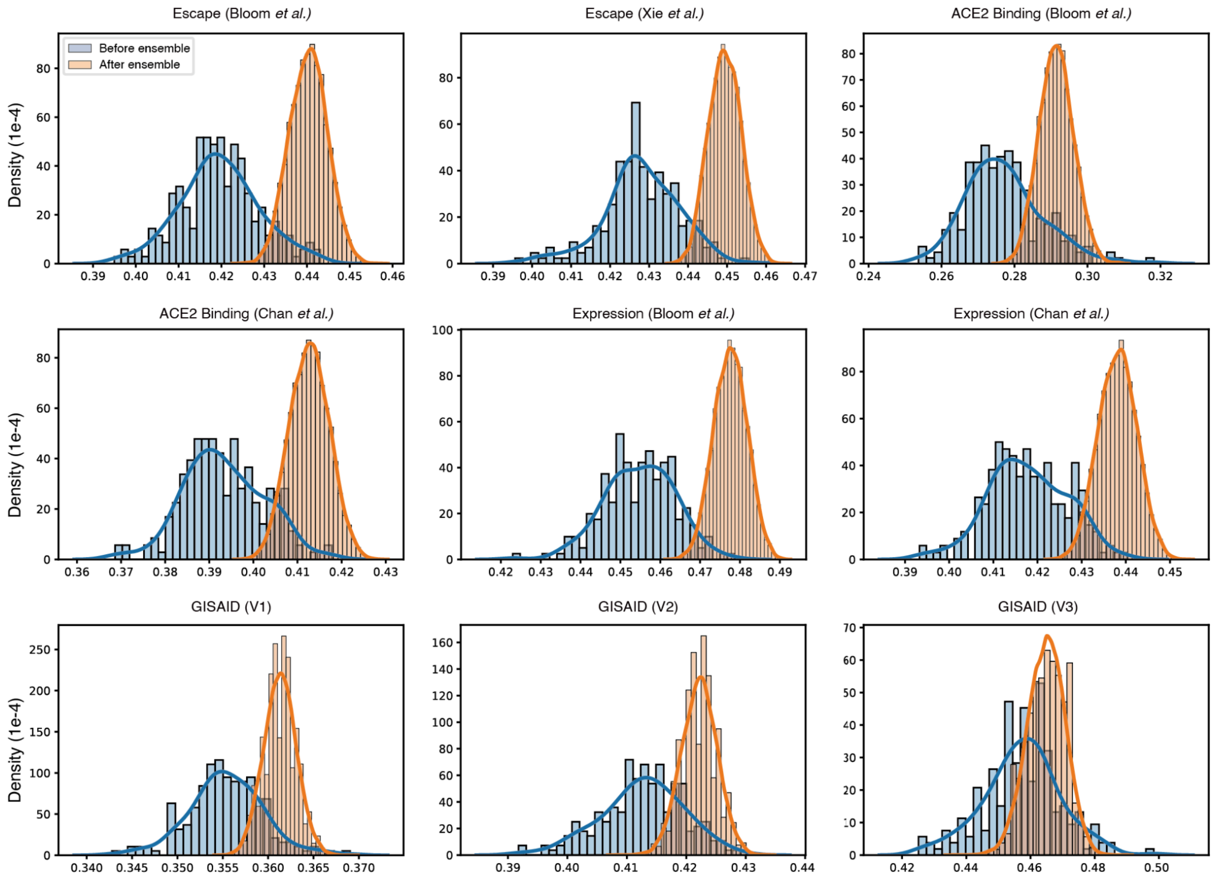


**Fig . S2. Distributional comparison of single models versus a 10-model ensemble across nine evaluation panels.** Blue histograms and density curves summarize the performance of individual models; orange histograms and curves show the corresponding 10-model ensembled predictions. For the six panels in the top two rows, performance is quantified by Spearman's rank correlation between model predictions and experimental measurements. For the three GISAID panels in the bottom row, the metric is the predicted ratio: the fraction of important pandemic sequences captured by the models' top-ranked predictions, evaluated at increasingly important thresholds from left to right (>10, >100, and >1000). Across panels, the ensemble distributions are consistently shifted to the right and are narrower than those of single models, indicating higher median performance and reduced variance.

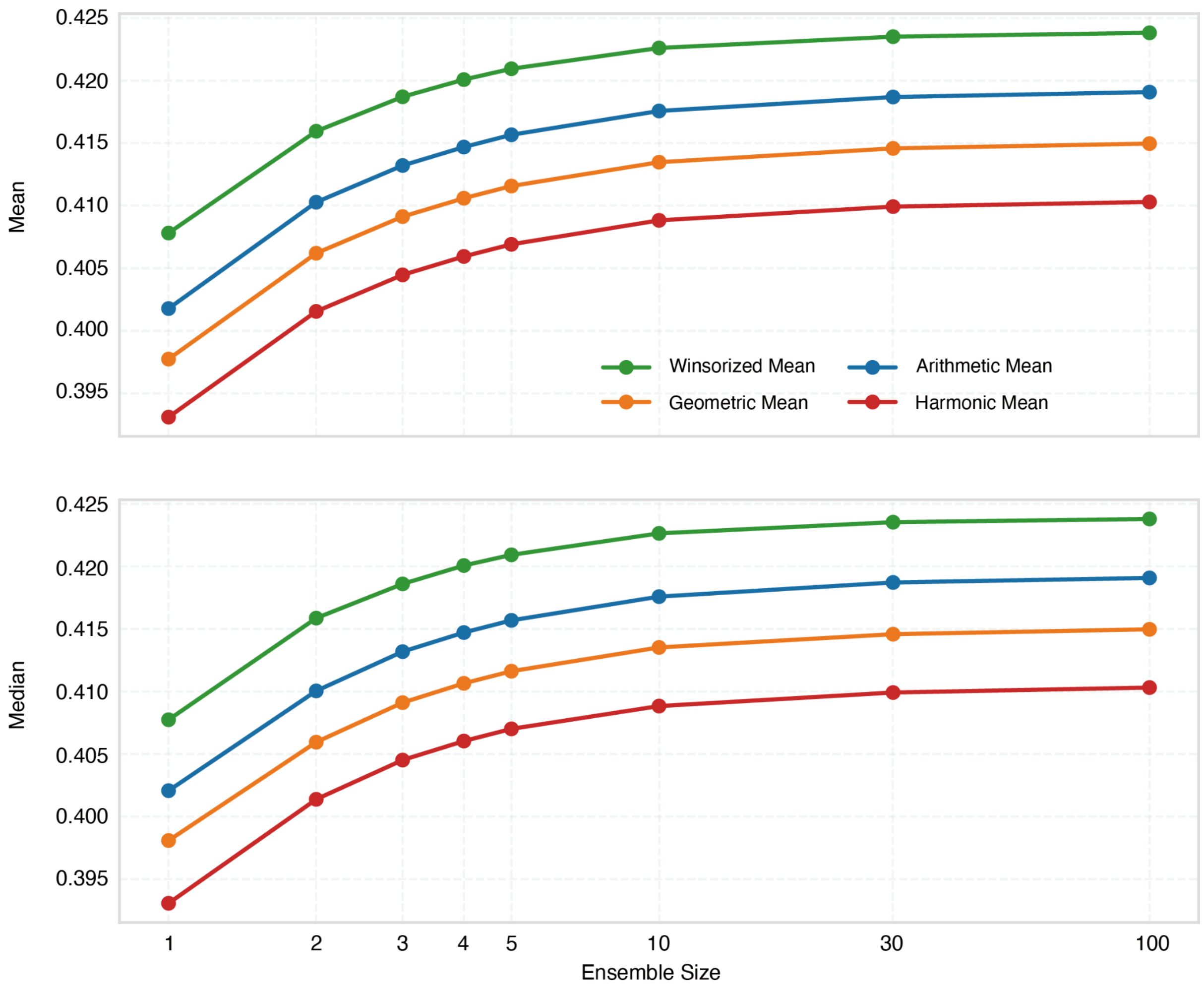


**Fig. S3. Accuracy effect of ensemble size on aggregate score across versions.** Top: Mean aggregate scores versus ensemble size (log scale). All four aggregate variants (v1–v4) improve monotonically with larger ensembles. v1 consistently attains the highest mean, followed by v2, v3, and v4. Bottom: Median aggregate scores versus ensemble size (log scale). The median exhibits the same monotonic trend and ranking as the mean. Improvements are most pronounced when increasing from size 1 to 5–10 and plateau thereafter.

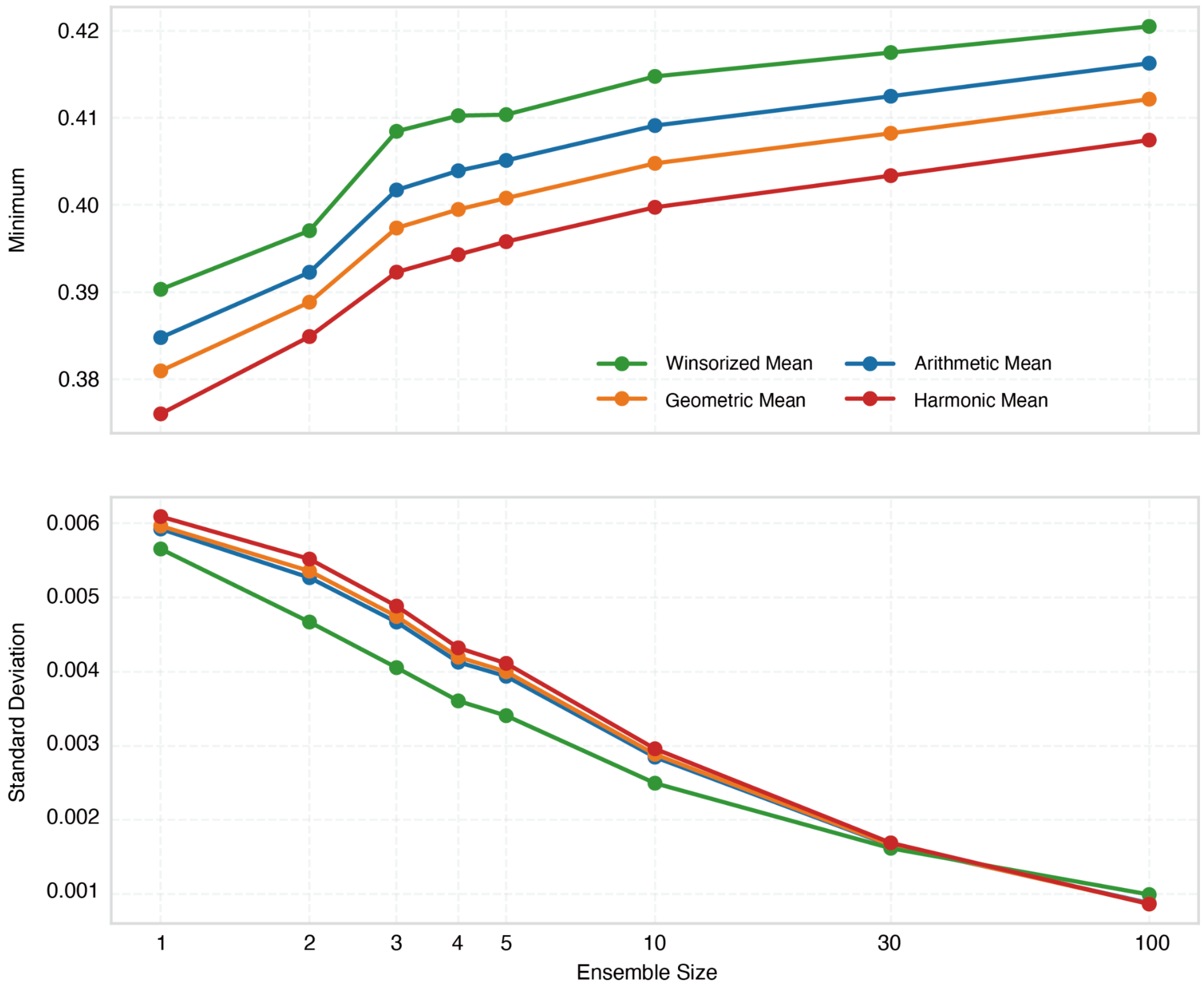


**Fig. S4. Robustness effect of ensemble size on aggregate score across versions.** Top: Minimum aggregate scores versus ensemble size (log scale). The worst-case performance improves steadily as ensembles grow, with the largest gains from size 1 to 5–10 and diminishing returns thereafter. Bottom: Standard deviation of aggregate scores versus ensemble size (log scale). Variability decreases approximately monotonically with larger ensembles, indicating increased robustness. Importantly, the standard deviation is approximately linear with log(ensemble size), indicating that variance decays smoothly with ensemble size on a logarithmic scale.

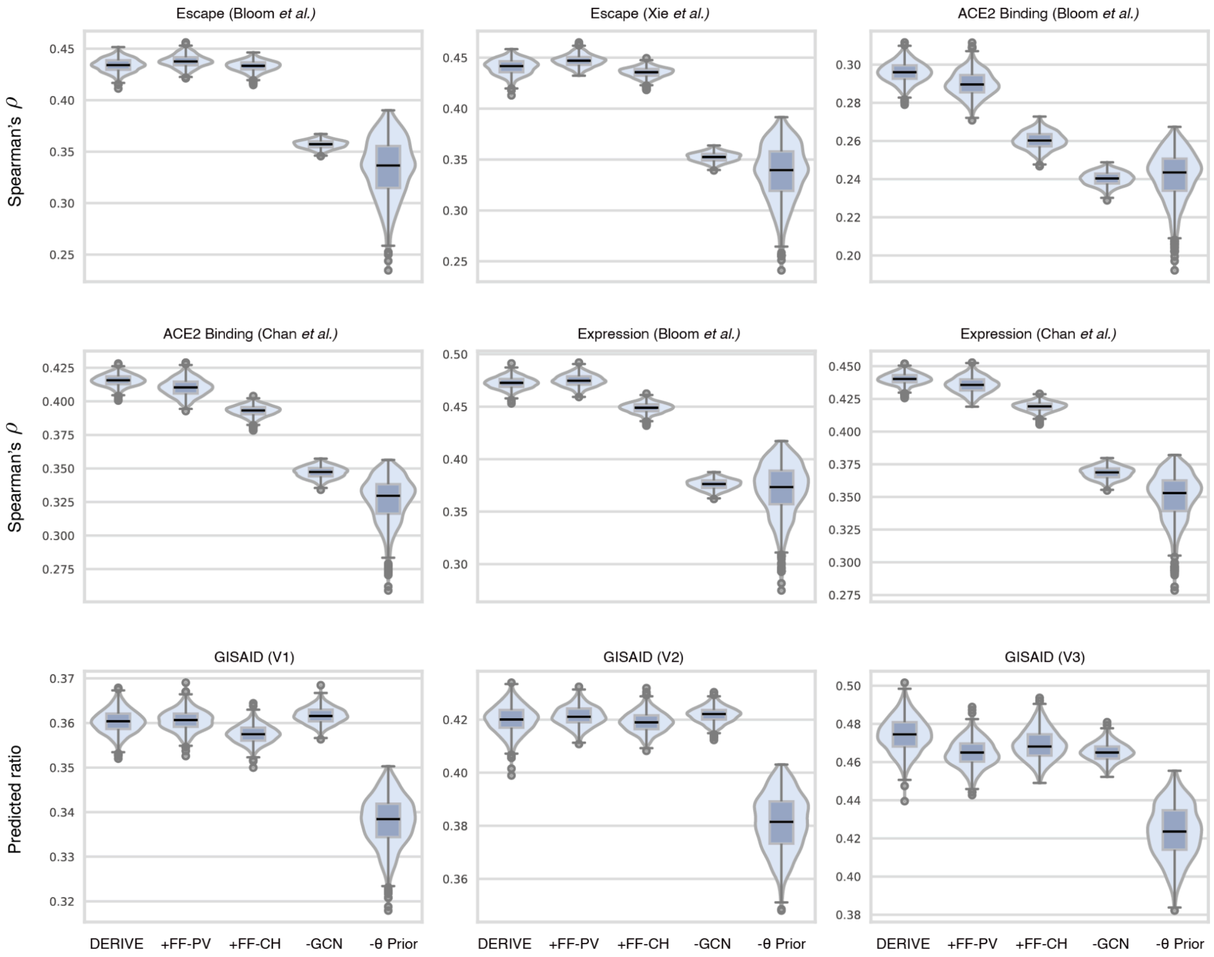


**Fig. S5. Ablation studies.** Here**,** Model ablation experiments investigated: "DERIVE" (the complete model); "+FF-PV" (substitutes polarity and side-chain volume attributes with parameters from coarse-grained force-field statistics); "+FF-CH" (replaces charge and hydrophobicity attributes with features derived from coarse-grained force-field statistics); "-GCN" (removes the GCN module and its associated augmentation); and "-θ prior" (eliminates parameter priors, resulting in a non-Bayesian VAE architecture). Performance metrics are defined as follows: for the six panels in the top two rows, the key metric is the Spearman's rank correlation between model predictions and experimental measurements. For the three panels in the bottom row (based on GISAID data), performance is given by the "predicted ratio": the fraction of significant pandemic sequences captured within the models' top-ranked predictions. This ratio was evaluated at thresholds of increasing significance from left to right (>10, >100, and >1000 sequences).
